# Improved RAD51 binders through motif shuffling based on the modularity of BRC repeats

**DOI:** 10.1101/2020.05.14.097071

**Authors:** Laurens H. Lindenburg, Teodors Pantelejevs, Fabrice Gielen, Pedro Zuazua-Villar, Maren Butz, Eric Rees, Clemens F. Kaminski, Jessica A. Downs, Marko Hyvönen, Florian Hollfelder

## Abstract

Exchanges of protein sequence modules support leaps in function unavailable through point mutations during evolution. Here we study the role of the two RAD51-interacting modules within the eight binding BRC repeats of BRCA2. We created 64 chimeric repeats by shuffling these modules and measured their binding to RAD51. We found that certain shuffled repeats were stronger than any of the natural repeats, suggesting balancing of relative properties in BRC repeats. Surprisingly, the contribution from the two modules was poorly correlated with affinities of natural repeats, with weak BRC8 repeat containing the most effective N-terminal module. The binding of the strongest chimera, BRC8-2, to RAD51 was improved by −2.44 kCal/mol compared to the strongest natural repeat, BRC4. Crystal structure of RAD51:BRC8-2 complex shows an improved interface fit and an extended β-hairpin in this repeat. BRC8-2 was shown to function in human cells, preventing the formation of nuclear foci after ionizing radiation.

## Introduction

The daunting combinatorial diversity arising from simultaneous mutation of all amino acid positions even in small proteins (leading e.g. to 10^39^ variants of a 30-mer) renders exploration of such sequence space futile. It is therefore attractive to view proteins not only as combinations of variant amino acids, but as combinations of exchangeable segments, because the combinatorial diversity is dramatically reduced if such ‘modules’ are instead recombined^1–6^. Understanding protein modularity at different length scales is key to elucidating natural protein evolution and to achieving full control in protein design and engineering. Shuffling is an empirically proven approach used in directed protein evolution^7–9^ and has been rationalized by the automated determination of minimally folded domains for shuffling, for example using the SCHEMA algorithm^10,11^. The exon-intron architecture of eukaryotic genes may in fact serve the modular evolution of proteins through *exon shuffling* – homologous recombination at introns to bring exons into novel combinations – and help bring about new protein functions^12–15^.

Here we address the relationship of modularity and function in a protein-protein interaction pair, RAD51:BRCA2, involved in DNA double-strand break repair. BRCA2 exerts a multitude of functions on RAD51 in the cell, such as localization, nucleofilament assembly and its depolymerization and has been aptly termed the ‘custodian’ of chromosomal numerical and structural integrity^16^. BRCA2 is a 3418 amino acid protein whose central part, exon 11, contains eight conserved repeats (referred to as ‘BRC repeats’ followed by the number 1 to 8, each consisting of around 35 residues^17^; Fig. 1a).

**Fig. 1.**
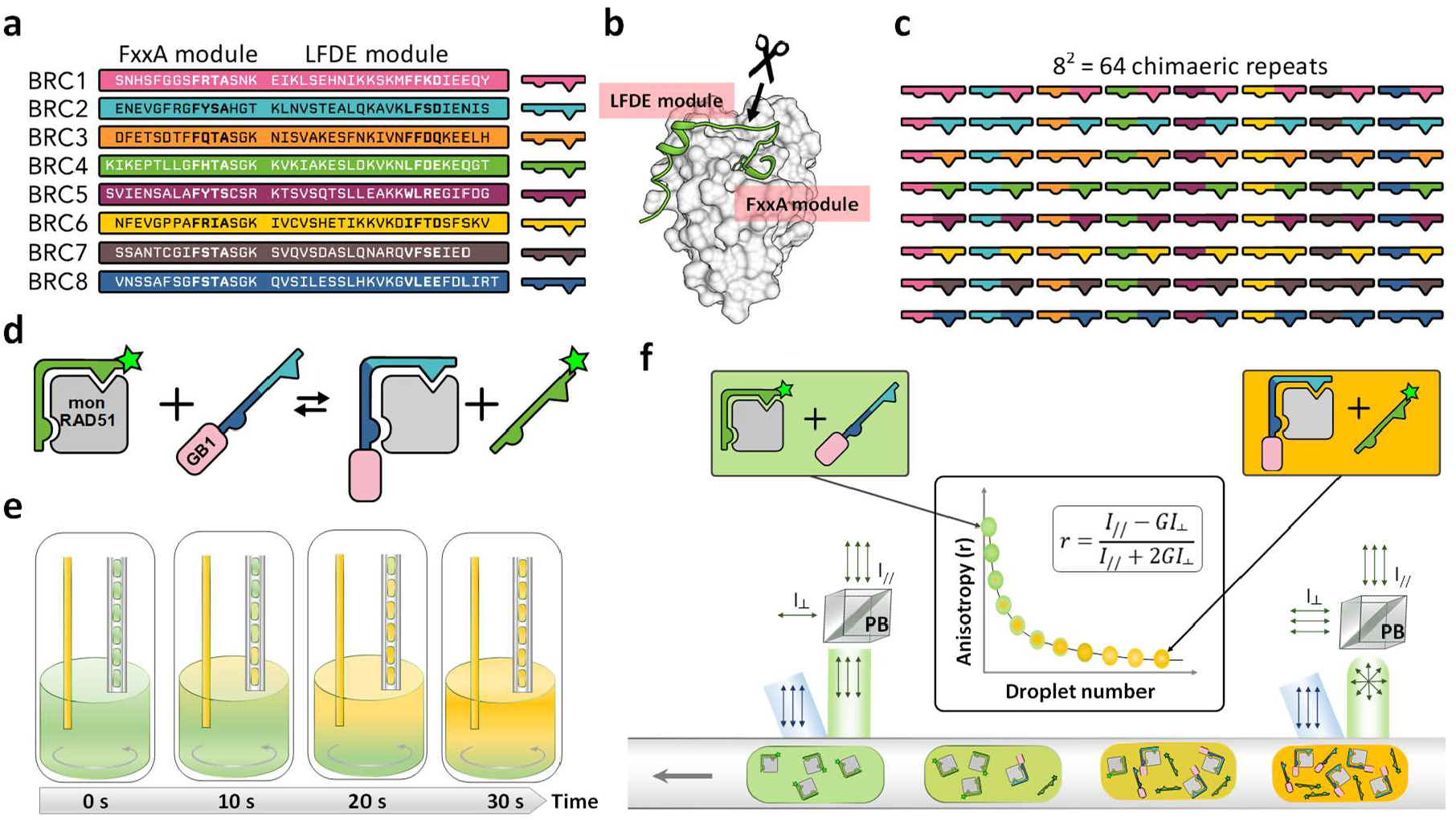
Shuffling of the binding modules comprising the eight RAD51-binding repeats in BRCA2. **a** The eight human BRC repeats with FxxA and LFDE motifs in bold. **b** The crystal structure of RAD51:BRC4 peptide complex (PDB ID: 1n0w). The arrow with scissors indicates the crossover point between the FxxA and LFDE modules used in this this study. **c** Schematic representation of the 56 chimeric and eight natural repeats resulting from the shuffling around the crossover point described in **b. d** Schematic representation of the competition assay used for the affinity determination of the BRC repeat shuffle set for monomeric RAD51 (mon RAD51) by fluorescence anisotropy. GB1-BRC recombinant peptide fusions were titrated into a complex of monomeric RAD51 and BRC4^fl^ (a fluorescein-labeled BRC4 synthetic peptide), so that the *K*_d_ of the recombinant peptide could be calculated using the known *K*_d_ of BRC4^fl^ ^27,28^. **e** Titration of GB1-BRC peptides using droplets-on-demand: microdroplets containing an increasing concentration of fusion peptides were produced from a well of a 384 well plate by a micro-capillary aspiration technique. During 30 seconds, a stock solution of GB1-BRC peptide was injected in the well so that its concentration rose from 0 to 50% of stock concentration (left capillary). Simultaneously, droplets encapsulating the changing contents of the well were generated (right capillary). Flow directions are represented by gray arrows. Mixing was performed by a magnetic stir bar (grey circling arrow). The concentration of monomeric RAD51 and BRC4^fl^ was kept constant throughout the titration. **f** The titration droplets were measured for their average fluorescence anisotropy using a fluorescence anisotropy imaging microscope. Upon travelling above the imaging area, linearly polarized light (in blue) excited the fluorescein tags of the BRC4^fl^ peptides, either bound to monomeric RAD51 (larger hydrodynamic volume, resulting in high anisotropy) or freely tumbling after competition with GB1-BRC peptides (lower hydrodynamic volume, resulting in low anisotropy). Fluorescence emission intensity signals (in green) recorded for parallel and perpendicular polarizations enabled the quantification of average anisotropy for every droplet of the titration sequence. A G factor was calculated to correct for differences in detector sensitivities in the parallel and perpendicular channels. PB=Polarizing Beamsplitter.

The crystal structure of BRC4 in complex with RAD51^18^, together with structural modeling and biochemical experiments, revealed the existence of two distinct parts in the BRC repeats that interact with RAD51^19^. The first of these, known as the ‘FxxA module’, forms a β-hairpin structure and binds RAD51 with a Phe and Ala in two small binding pockets of the ATPase domain. The C-terminal part of the BRC repeat, with a conserved LFDE motif, interacts with the distal part of the ATPase domain in an α-helical conformation. In doing so, the BRC repeats directly compete with another FxxA module located in RAD51 itself (with the sequence FTTA), on an oligomerization epitope between RAD51’s C-terminal ATPase and N-terminal DNA-binding domains. The BRC4 repeat peptide has been shown to cause dissociation of RAD51 oligomers and conditional expression of the repeat in breast cancer cells disrupts the RAD51:BRCA2 interaction and sensitizes them to radiation treatment^20^. In isolation, the FxxA module makes a relatively weak contribution to the binding: a 4-residue FHTA peptide, representing the FxxA hotspot from the corresponding module in BRC4, bound RAD51-surrogate HumRadA2 with a *K*_d_ of 290 μM^21^. Even the entire FxxA module – that is the FxxA hotspot and the surrounding residues – is not a strong binder of RAD51; about 500 μM of a 17-residue FHTA-containing peptide (the N-terminal half of the BRC4 repeat) was required to effect full disruption of the RAD51:BRC4 interaction in a competitive ELISA assay^19^. It is the C-terminal LFDE module that ensures significantly enhanced affinities are achieved. This second module binds to a groove on another surface of the RAD51 ATPase domain^18^. Although the phylogeny of the BRC repeats remains to be fully elucidated, it is thought that the emergence of the BRCA2 repeats predates the emergence of the mammalian class^22^ and perhaps even the divergence of birds and mammals 230-300 million years ago^23^. As the eight repeats found in the BRCA2 protein all occur on the same exon^22^, throughout their evolutionary history, these repeats would never have been subject to natural exon shuffling.

Recognizing this modularity, we probe the idea that functional sophistication is brought about by combination of these relatively simple peptide building blocks, by testing whether rearranged BRC repeats can lead to functional chimeras that interact with RAD51. Would artificial shuffling of repeat modules engender affinity maturation by bringing together the most binding-proficient FxxA and LFDE modules and could such chimeric peptides give us improved insight into module-specific contributions to RAD51 binding? Our objective was to explicitly explore the recombination of entire natural repeats in the creation of new functional proteins, in order to demonstrate the role of modularity in functional adaptation. We discovered that the natural, ‘parental’, combination of modules often turned out to be suboptimal for RAD51 binding, and that upon decoupling of natural BRC module combinations, more potent RAD51 binders could be obtained.

## Results

### Systematic shuffling of the two binding modules of the eight BRC repeats and evaluation of the 64 resulting chimeras in a microfluidic fluorescence anisotropy assay

To be able to shuffle the two modules of the eight BRC repeats found in BRCA2 that bind RAD51 (Fig. 1a), a crossover point was defined immediately at the C-terminus of the hairpin structure found in the FxxA module (Fig. 1b), as suggested by the RAD51:BRC4 crystal structure^18^. The resulting 64 variant BRC peptides (Fig. 1c) were cloned through an oligonucleotide cassette method (Supplementary Fig. 1 & Supplementary Table 1), as C-terminal fusions to the GB1 domain from protein G (Supplementary Fig. 2; we will refer to shuffled variants by two digits denoting the identity of the N-terminal FxxA and the C-terminal LFDE module, respectively, e.g. BRC2-4 is a peptide with the FxxA module from BRC2 and the LFDE module from BRC4). The GB1 domain does not interfere with the BRC-repeat interaction: an isothermal titration calorimetry (ITC) affinity measurement was carried out to confirm that the BRC4 peptide, upon fusion to the GB1 C-terminus, maintained its ability to bind HumRadA22 (a faithful yet monomeric model of RAD51 and for simplicity we will refer to this as monomeric RAD51^24^; Supplementary Figs. 5, 6 and accompanying Supplementary Text 3) and matched affinities previously measured for BRC4 peptide^24–26^.

The 64 different variant peptide repeats were assayed in a competition assay with a fluorescently labelled BRC4 repeat (BRC4^fl^-peptide) as the tracer (Fig. 1d). We employed a microfluidic setup to facilitate measurement of affinities to minimize the sample quantities^27^: dose-response curves were set up by coupling ‘droplet-on-demand’ formation (Fig. 1e) and a microfluidic fluorescence anisotropy imaging platform (Fig. 1f), so that every measurement only required nanoliter volumes and minimal protein amounts.

Although the droplet-on-demand method supported procurement of measurements over at least two orders of magnitude in concentration of titrant, we expected even larger differences in affinity between the different chimeras. Consequently, the BRC repeats were diluted to an appropriate concentration (as established by an initial, single concentration-point screen, Supplementary Table 3) for acquisition of a dose-response profile in microdroplets, representing individual titrations of the chimeras (Supplementary Fig. 7). The small amounts of peptides necessary to carry out droplet assays, made it possible to screen close to saturating conditions (with the exception of the poorest binders), resulting in good quality data which could be fit to a competitive binding model (Fig. 2a) to derive dissociation constants for all 64 variants (Fig. 2b).

### Shuffling leads to BRC peptide binders with enhanced affinity over wildtype

The 64 measured *K*_d_ values for BRC peptide binding to monomeric RAD51 spanned a range of three orders of magnitude from 11 μM (BRC7-8) to 6 nM (BRC8-2). To the best of our knowledge, most of the shuffled variants had never been measured before (except BRC4-5 and BRC5-4^19^), so it is useful to compare the values we measured here for the natural repeats (which can be read as a diagonal from the top left corner to bottom right corner in Fig. 2b) to previously reported values. First, using this competitive fluorescence anisotropy assay, BRC4 was found to have a *K*_d_ of 38 nM for monomeric RAD51, slightly higher than the value previously determined by direct titration by fluorescence anisotropy (12 nM^27^) or by ITC in this study (11 nM, Supplementary Fig. 5b, g). These values fall within the range of previously measured affinities (6.2-64 nM) for this protein-protein interaction^24^. Also, BRC4 was found to be the tightest binder of all the natural repeats, in agreement with previous studies^29,30^. The repeats 1, 2, 3 and 4 displayed higher affinity (median *K*_d_ 198 nM) than repeats 5, 6, 7 and 8 (median *K*_d_ 1394 nM). This observation is consistent with previous reports that BRC repeats 1 to 4 have higher affinity than repeats 5 to 8 for uncomplexed RAD51^30^. Beyond this broad analysis, more detailed comparisons (including the absolute *K*_d_ values) to previous studies are of limited value as there are important differences in the assays employed and the binders titrated against, e.g. full length RAD51^30,31^ or truncated RAD51 consisting of the catalytic domain only^29^.

Having established that the affinity ranking for the natural repeats were consistent with previous reports, we next analyzed the affinities we found for the novel recombinant peptides. The combinations with the FxxA module of BRC5 were found to be the weakest binders, easily explained by the fact that the conserved alanine in the FxxA module of BRC5 is replaced by a serine (FYTS), which has a hydrophilic side-chain that would not form favorable steric and electrostatic contacts with the compact Ala pocket. Interestingly, the recombinant peptide BRC4-5 was also found to be a relatively weak binder (*K*_d_ 2.1 μM), despite previous findings that this chimera displayed a stronger affinity19. This paradox is addressed below. Remarkably, a few chimeras containing the FxxA module from BRC8, a natural repeat from the ‘weak’ group of repeats 5 to 8, were found to be the strongest binders from the entire set of 64 variants (Fig. 2b) with BRC8-2 being the peptide with highest affinity with a *K*_d_ of 6 nM. This highlights the that shuffling can lead to the bringing together of elements that are in a non-optimal combination in nature.

**Fig. 1.**
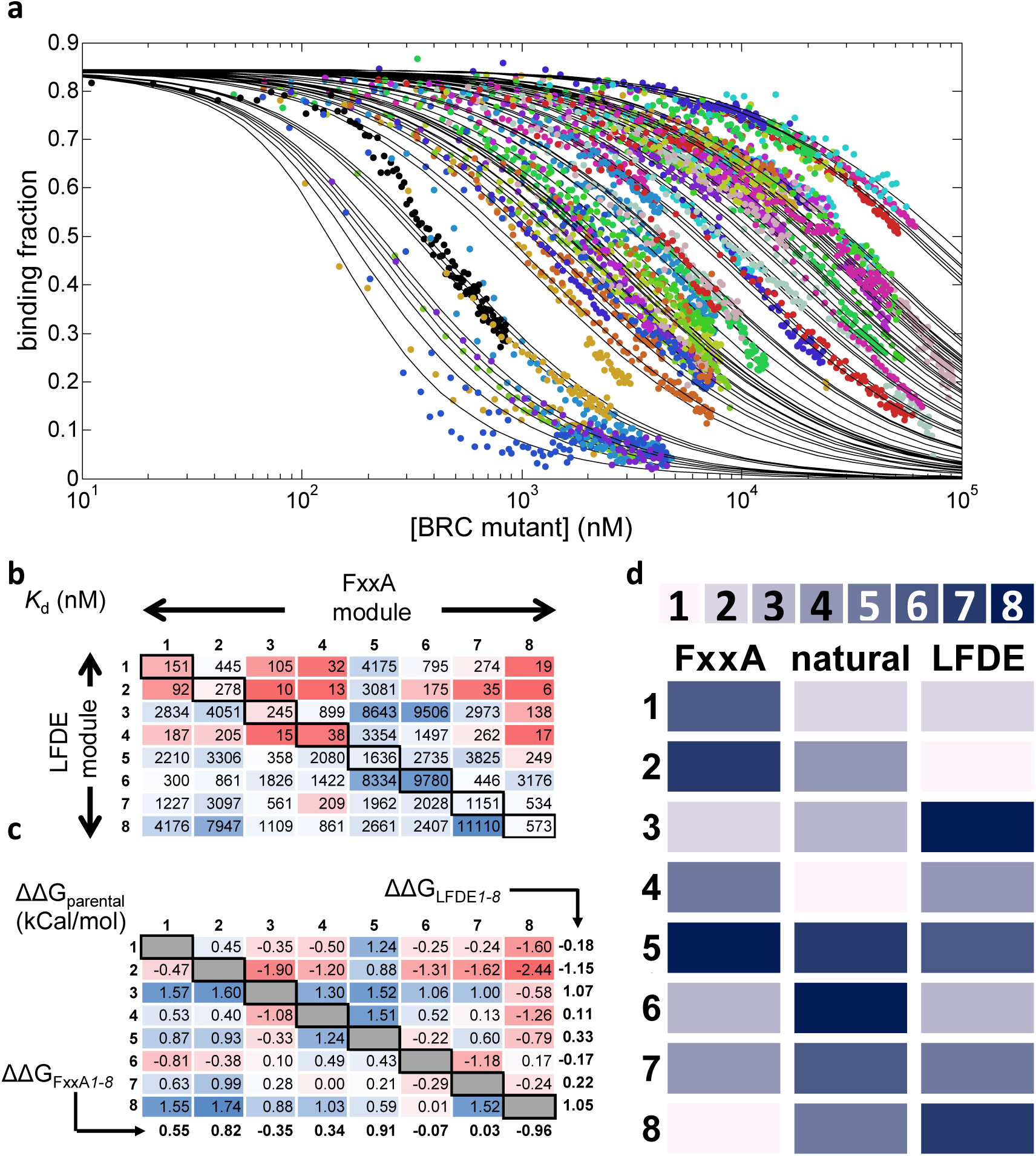
Affinity determination of 64 chimeric BRC4 repeats by a microfluidic droplet-on-demand system interfaced with fluorescence anisotropy detection. **a** Fraction of BRC4^fl^-peptide bound to monomeric RAD51 as a function of GB1-BRC peptide chimera concentration, measured using the droplet-on-demand anisotropy competition assay. Measurement conditions were 100 nM BRC4^fl^, 150 nM monomeric RAD51 in a buffer of 20 mM CHES (pH 9.5), 100 mM NaCl, 1 mM EDTA, at 20 °C. Note the starting binding fraction of 0.85 that is calculated from the affinity of the BRC4^fl^ peptide for monomeric RAD51 and the initial concentrations used^28^. **b** *K*_d_ values (in nM) determined for all 64 BRCA2 peptide chimeras using data in **a. c** Analysis of the effect of recombination, expressed as the difference in ΔG (Supplementary Table 4) of each shuffled variant relative to the average of the two natural parental combinations (ΔΔG_parental_, in kCal/mol) for each variant. The parental combinations are depicted in grey (by definition, their ΔΔG_parental_ is always zero). The values indicated below each column and next to each row represent the average ΔΔG_parental_ value for FxxA and LFDE modules from each repeat, respectively and are referred to as ΔΔG_FxxA*1-8*_ and ΔΔG_LFDE*1-8*_, respectively. **d** Binding rank order of the parental BRC repeats (central column) and the individual FxxA (left-hand column) and LFDE (right-hand column) modules comprising the repeats, indicated by intensity (low intensity = top binder, high intensity = poor binder, as indicated by scale bar). The contributions to binding of the 8 different repeat-derived modules are calculated using their ΔΔG_LFDE*1-8*_ and ΔΔG_FxxA*1-8*_ values.

### Discerning module-specific contributions to binding affinity and the effect of the crossover point placement

To allow comparisons across repeats and modules, the dissociation constants were expressed in units of Gibbs free energy (ΔG, Supplementary Table 4). The effect of shuffling was quantified by expressing each of the 56 novel, unexplored combinations in terms of relative change vis-à-vis their two parental repeats, e.g. the two parents of BRC1-2 are BRC1 and BRC2 (Fig. 2c and Supplementary Information Text 5.1). A positive value for ΔΔG_parental_ – indicating that the product of the shuffling was detrimental to the binding function – was observed in 33 out 56 peptides. Also, the average ΔΔG_parental_ for all 56 shuffled repeats was 0.16 kCal/mol, indicating that shuffling had a net-detrimental effect on binding function. Nevertheless, 23 out of these 56 repeats had negative ΔΔG_parental_ values and thus represented variants that were improved over the average of their parents. We asked whether the identity of the FxxA module affected binding to monomeric RAD51 more strongly than the identity of the LFDE module. At a first approximation, the FxxA module might appear to have dominated the interaction as proven by the fact that any combination with imperfect FxxA module of BRC5 (containing the stretch of residues Ser1662^BRC5^ to Arg1677^BRC5^ and in which the alanine of the FxxA motif is replaced by a serine) resulted in exceedingly weak interactions (ΔΔG_FxxA*5*_ = 0.91 kCal/mol) (Fig. 2c). The LFDE modules from BRC3 and BRC8 were found to cause the most significant reduction to binding in each of their respective seven recombinant peptides (ΔΔG_LFDE*3*_ = 1.07 kCal/mol; ΔΔG_LFDE*8*_ = 1.05 kCal/mol). As both the FxxA and LFDE modules in each repeat could thus make a significant contribution to binding, we considered whether the net contribution to binding was equally distributed within each repeat. In BRC repeat 5, both the FxxA and LFDE modules conspired to make a weak binder (ΔΔG_FxxA*5*_ = 0.91 kCal/mol; ΔΔG_LFDE*5*_ = 0.33 kCal/mol). By contrast, BRC repeat 8 could be considered ‘Janus-faced’, as it is composed of a net contributor (ΔΔG_FxxA*8*_ = −0.96 kCal/mol) and a net disruptor (ΔΔG_LFDE*8*_ = 1.05 kCal/mol) to binding. To a slightly lesser degree, BRC repeat 2 displayed the same contrast in intra-repeat properties, although in this natural repeat, the FxxA module was a net disruptor overall (ΔΔG_FxxA*2*_ = 0.82 kCal/mol), while the LFDE module was a net contributor (ΔΔG_LFDE*2*_ = −1.15 kCal/mol). This analysis is validated by the observation that BRC8-2, which combines the overall best FxxA module with overall best LFDE module, is the BRC peptide with the highest affinity of all, which we were able to cross validate by ITC measurements (Supplementary Fig. 5) and is also the most improved over its parental sequences (ΔΔG_parental_ = −2.44 kCal/mol). The depiction of the rank affinity ordering of individual modules by intensity, based on their ΔΔG_FxxA1-8_ or ΔΔG_LFDE1-8_ values, next to the rank order of the natural repeats’ affinities (Fig. 1d), highlighted that within the eight natural BRC repeats, binding function was not always equally distributed between modules. It is also interesting to note that the two modules of BRC4, the highest affinity natural repeat, are relatively poor contributors of affinity and highly dependent on which module they pair with. In contrast, BRC6, the natural repeat with the lowest affinity in our analysis, contains modules that, in combination with other modules, contribute positively to affinity.

The observation of a ΔΔG_parental_ of 1.24 kCal/mol (Fig. 2c) for BRC4-5 represented a notable discrepancy to previous data, obtained by competitive ELISA with synthetic chimeric peptides BRC4-5 & 5-4 ^19^. As expected, in agreement with our findings, BRC5-4 turned out to be a weak binder, due to the lack of conservation in repeat 5’s FxxA module. However, Rajendra and Venkitaraman ^19^ found BRC4-5 to be a *stronger* binder than the natural BRC4, whereas we found BRC4-5 to bind monomeric RAD51 with 55-fold *lower* affinity than BRC4. What could explain this? Apart from the obvious difference in the assays (heterogeneous ELISA-based assay vs homogeneous polarization-based assay), the main remaining difference is the cutoff between the end of the FxxA module and the start of the LFDE module in the shuffled peptide. While Rajendra & Venkitaraman ^19^ defined Lys1533^BRC4^ as the last residue of the FxxA module and Ile1534^BRC4^ as the first of the LFDE module, our peptides were based on the cut-off point occurring between Lys1530^BRC4^ and Lys1531^BRC4^. The cutoff is arbitrary but resulted in our chimeric BRC4-5 repeat bearing two mutations compared to the other study, Val1532^BRC4^ → Thr1679^BRC5^ and Lys1533^BRC4^→ Ser1680^BRC5^. Retrospective analysis using SCHEMA computational algorithm to identify the optimal cross-over points in a protein sequence ^10^ found a cross-over point between Ile1534^BRC4^ and Ala1535^BRC4^ to be optimal (Supplementary Fig. 8 & Supplementary Text 5.2). A cut-off point more distant from the FxxA hairpin results in fewer pairwise interactions (i.e., between residues from the N-terminal half with residues from the C-terminal half) being broken upon shuffling. Furthermore, we found that the deletion of Lys1530^BRC4^, located at our chosen cross-over point, resulted in a significant loss of affinity (the *K*_d_ shifted almost 200-fold, from 21 nM to 4.1 µM; Supplementary Fig. 9 & Supplementary text 5.3). Thus, subtle differences in the linker region can lead to dramatic differences in affinity, explaining the effect of the exact placement of shuffle cut-off points to the relative contribution of the two modules.

### BRC8-2 forms a more extensive β-hairpin and has improved helicity compared to BRC4

To gain structural insight into the increased affinity of BRC8-2 for RAD51, we determined a 1.95 Å crystal structure of the monomeric RAD51:BRC8-2 complex (PDB ID: 6HQU). There are eight complexes in the asymmetric unit of these crystals, all of which are very similar to each other with an average RMSD of 0.664 Å for 198 Cα atoms of RAD51. Bound BRC8-2 peptide is visible in seven of the eight RAD51 molecules, in essentially identical conformation in all complexes. Representative electron densities before the BRC8-2 has been modelled and after final refinement are provided in Supplementary Fig. 10. The best-defined complex (chains B and J) has been used in the subsequent analyses.

Comparison of the refined structure for monomeric RAD51:BRC8-2 with that of the RAD51:BRC4 complex (PDB: 1n0w; 18) shows a similar overall topology (Fig. 3a). Phe2058^BRC8^ and Ala2061^BRC8^ of the BRC8 FxxA module form identical contacts to those seen between BRC4 and RAD51. C-terminal to Ala2061^BRC8^, the peptide forms a β-hairpin that extends the central β-sheet of monomeric RAD51 in an inter-molecular fashion, reminiscent of the RAD51:BRC4 complex^18,32^. Five residues at the C-terminal ends of the BRC8 and BRC4 FxxA modules are identical in sequence (TASGK) and both hairpins are stabilized by the hydroxyl groups of Thr2060^BRC8^/Thr1526^BRC4^ forming hydrogen bonds with the backbone amine of Lys2064^BRC8^/Lys1530^BRC4^ and the hydroxyl of Ser2062^BRC8^/Ser1528^BRC4^ (Fig. 2b). In the RAD51:BRC4 complex, the C-terminal LFDE module forms a ten-residue α-helix that interacts with RAD51 through a shallow interface using a mixture of hydrophobic and polar contacts. In the BRC2 LFDE module (with sequence LFSD) Leu1240^BRC2^ and Phe1241^BRC2^ bind the same hydrophobic interface that BRC4 interacts with and Asp1243^BRC2^ interacts with a nearby Arg270^monomeric RAD51^ as seen in BRC4 (Fig. 3c). Contact areas calculated for the BRC4 and BRC8-2 complexes are very similar, 1042 and 942 Å^2^, respectively, consistent with the previously noted weak correlation between buried surface area and binding affinity^33^.

The most significant difference between the BRC4 and BRC8-2 peptides is the extent of the intra-molecular hydrogen-bonding network that forms the β-hairpin in the FxxA module. In BRC8-2, the β-hairpin is significantly extended, with its N-terminal end, before Phe2058^BRC8^, folding back towards the rest of the peptide (Fig. 2b). The hairpin extends a total of 19 amino acids, from Ser2053^BRC8^ to Thr1231^BRC2^ and is significantly longer than the nine-residue hairpin in BRC4. Formation of the extended hairpin is enabled by Ser2056^BRC8^, whose side chain fits tightly between the two anti-parallel strands of the peptide and the surface of RAD51, hydrogen bonding with the carbonyl of Leu1227^BRC2^ and the NH of Phe2058^BRC8^. This allows the peptide to fold back on itself and to form the extended intra-molecular H-bonding network (Fig. 3b). BRC4 has a larger, hydrophobic Leu1522 in the equivalent position of Ser2056^BRC8^, which cannot satisfy the steric and electrostatic requirements of the topology we observe in BRC8-2, forcing the N-terminus of the peptide to point away from the β-hairpin and the rest of the peptide. Interestingly, the FxxA module of BRC3 contains a threonine in the position equivalent to Ser2056 and has the second highest ΔΔG_FxxA1-8_ (ΔΔG_FxxA3_ = −0.35 kCal/mol). BRC3 is likely to form a similar extended hairpin to BRC8, with its threonine forming equivalent hydrogen bonds to Ser2056. To examine the contribution of Ser2056 to binding, we designed a mutant repeat BRC8-2(S2056A). Its affinity for monomeric RAD51 was measured as a *K*_d_ of 5 nM, i.e. an order of magnitude lower affinity than that measured for untagged BRC8-2, confirming the significance of Ser2056 for binding (Supplementary Fig. 5e,f).

The binding modes of the LFDE module of BRC8-2 shows also subtle differences compared to BRC4 (Fig. 2c). In BRC4, the bulky side chain of interface-forming Val1542^BRC4^ pushes the peptide away from RAD51, forming an outward-facing bulge and disrupting the optimal helical geometry of the peptide. The equivalent residue in BRC8-2 is smaller, Ala1237^BRC2^, and allows the α-helix to form a closer interaction with monomeric RAD51 and to retain a more regular helical geometry. Thus, the increased binding affinity of the BRC8-2 repeat for monomeric RAD51 appears to result from the extended hydrogen-bonding network of the FxxA module of BRC8 and the improved packing and helical geometry of the LFDE module from BRC2.

**Fig. 2.**
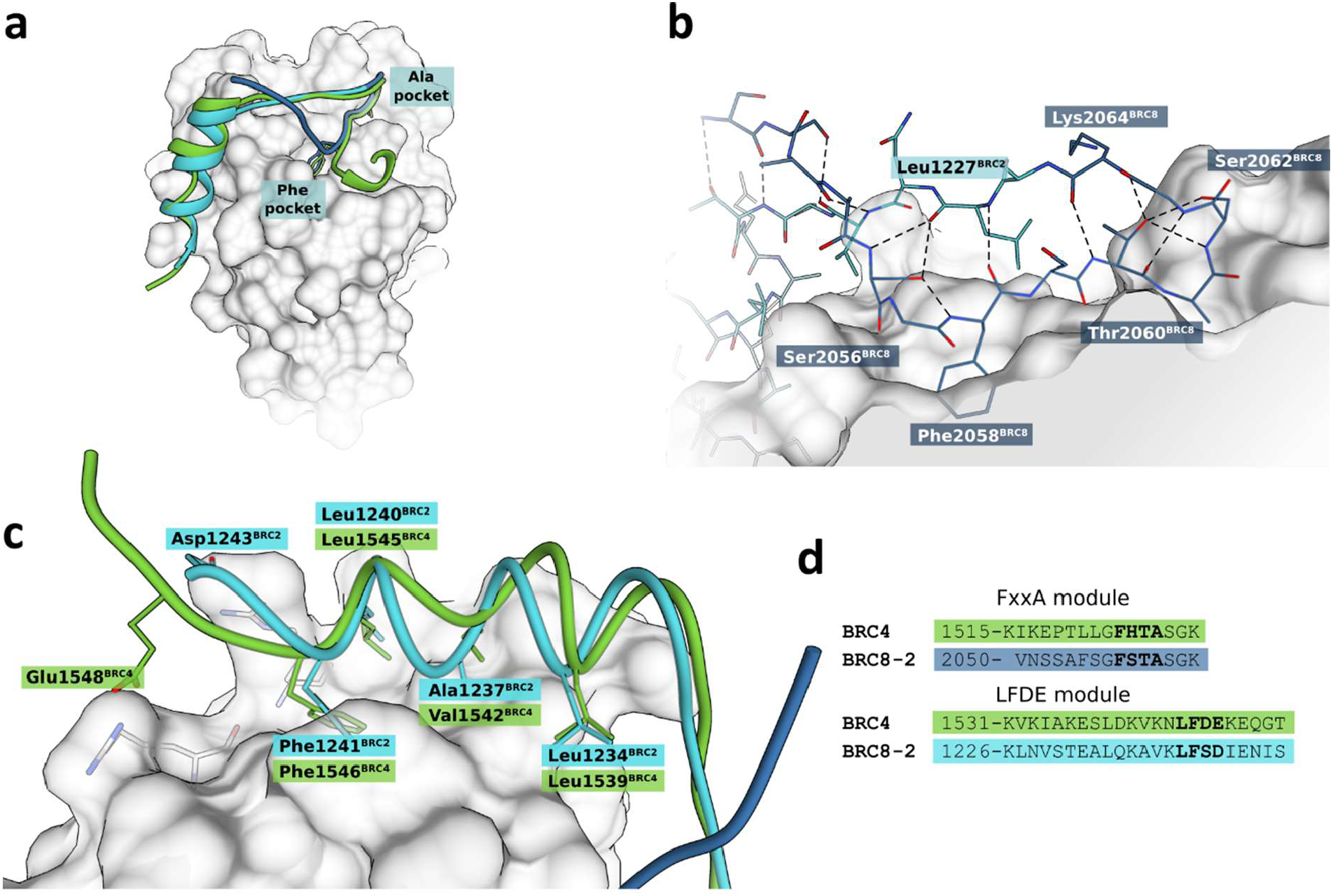
Crystal structure of the monomeric RAD51:BRC8-2 complex compared with RAD51:BRC4. BRC8-2 is depicted in dark and light blue, corresponding to BRC8 and BRC2 sequences, respectively. BRC4 is shown in green. Peptides were superimposed by aligning the structures of their respective protein binding partners. Monomeric RAD51 is represented by a grey surface. Selected residues of the monomeric RAD51 are depicted in grey. **a** Overall topologies of the two peptides, with the Phe and Ala pockets of the FxxA site shown. **b** Hydrogen-bonding network of the BRC8-2 β-hairpin. **c** LFDE interface with side chains of crucial residues depicted. **d** Sequence alignment of BRC4 and BRC8-2 FxxA and LFDE modules.

### BRC8-2 can disrupt RAD51 foci formation in cell-based experiments

To validate the utility of improved binding of BRC8-2 binding to RAD51 for biological intervention, we investigated the ability of this peptide to disrupt RAD51 function in human cells. Following treatment with ionizing radiation (IR), RAD51 translocates to the sites of DNA damage and forms foci that are visible by immunofluorescence. The formation of such foci is dependent on BRCA2^34^, and it has previously been shown that foci formation can be disrupted by expression of native BRC repeats^35,36^.

To determine whether BRC8-2 impaired RAD51 foci formation, the peptide was fused to a green fluorescent protein (GFP) containing a nuclear localization signal (NLS) and transfected into U2OS osteosarcoma cells. Cells were transfected at the same time with the parental GFP-NLS construct and RAD51 foci formation was monitored in GFP positive cells (Fig. 4a). As expected, the control GFP cells showed an increase in the median number of RAD51 foci after exposure to ionizing radiation (IR, 3 Gy) (Fig. 4b, c). In addition, a small number of foci were present in the absence of IR, most likely reflecting HR events associated with replicative stress. In contrast, cells expressing GFP-BRC8-2 had fewer RAD51 foci prior to irradiation and following IR exposure (Fig. 4b, c). Because RAD51 foci formation is limited to the S and G2 phases of the cell cycle, we wanted to determine whether the reduction in foci formation was due to an increase in G1 phase cells in the GFP-BRC8-2 expressing cells. We therefore monitored the cell cycle profile of GFP expressing cells by FACS and found that the GFP-BRC8-2 expressing cells have a decrease in the proportion of G1 phase cells, indicating that the effect on RAD51 foci formation is not due to cell cycle alterations (Fig. 4d, e). These data suggest that BRC8-2 interferes with RAD51 foci formation in human cells through binding and sequestering RAD51 away from sites of DNA damage. In support of this, we also noted that the pan-nuclear signal of RAD51 in GFP-BRC8-2 expressing cells is greater than in the GFP control cells (Supplementary Fig. 11).

**Fig. 4.**
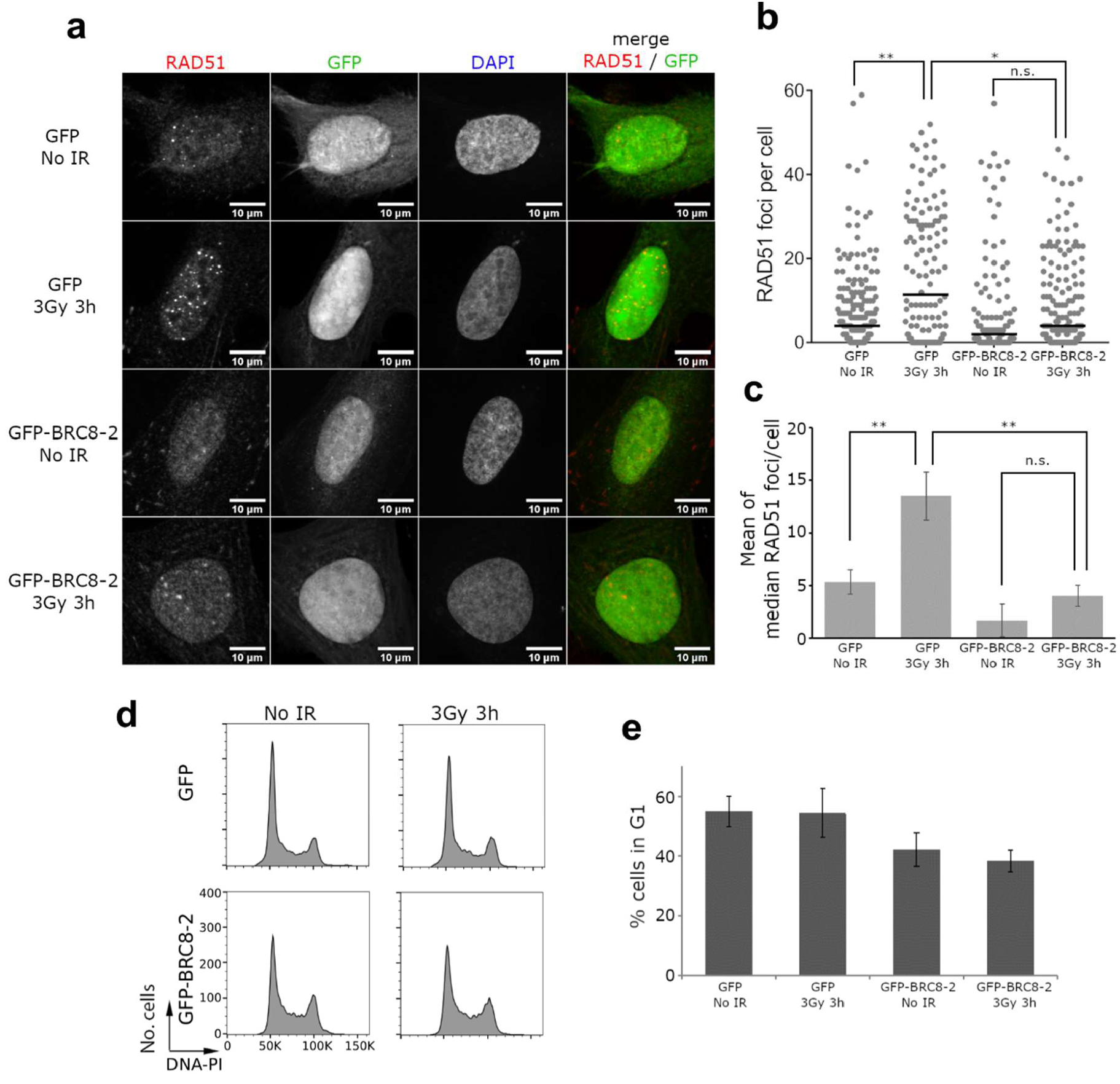
BRC8-2 impairs RAD51 foci formation in human U2OS cells. **a** Representative images of U2OS cells expressing empty GFP with a nuclear location signal (GFP) or GFP-BRC8-2 peptide (GFP-BRC8-2). Cells were monitored 3 hours after no treatment (No IR) or irradiation with 3 Gy (3Gy 3h) for GFP fluorescence or stained with RAD51 or DAPI as indicated. **b** Dot plot graph from one biological replicate plotting the number of RAD51 foci per GFP positive cell. Median values for each population are indicated with a bar. Outlier values were excluded from the graphical representation but included in the median calculation. More than 85 GFP positive cells were analyzed for each condition. Statistical analysis was done using Kruskal Wallis rank sum test followed by Dunn’s procedure for pairwise comparison (*p <0.005, **p<0.001, n.s.= not significant). **c** Bar graph showing the average median RAD51 foci per GFP positive cell from three independent biological experiments. Data are presented as mean ± SEM, n = 3 biological repeats. **p<0.001, n.s.= not significant, using ANOVA test (F(3,8)=32.03, p<0.0001) followed by Tukey’s method. **d** Representative cell cycle profiles from GFP positive cells transfected with GFP or GFP-BRC8-2. Cells were analyzed by FACS 3 hours after no treatment (No IR) or irradiation with 3 Gy (3Gy 3h). PI = propidium iodide. **e** Bar graph showing the percentage of cells in G1 phase. Data are the mean values from three independent biological experiments ± SD.

## Discussion

### Shuffling of BRC modules: anciently diverged parts meet

In this work we successfully exploited a droplet-on-demand platform to comprehensively survey the effect of shuffling of the BRC modules. This approach revealed novel BRC repeat combinations with unprecedented affinity for monomeric RAD51 and demonstrated *in cellulo* activity. Due to our systematic analysis of the contribution of each module to binding, as well as our crystallographic study of the monomeric RAD51:BRC8-2 complex, we contribute to the understanding of the structure-function relationship within the BRCA2:RAD51 complex, a highly dynamic and context-specific protein-protein interaction: although repeats 1 to 4 were found to bind *free* RAD51 more strongly than repeats 6 to 8, by us and others, repeats 5 to 8 were reported to have higher binding affinity for the RAD51:ssDNA complex than repeats 1-4^30,37^. Repeats 5-8 may also bind in concert to stimulate certain RAD51 functions^38^.

If combinations of modules achieve a wide range of affinities, their assembly context may matter (in addition to intrinsic effects of each module), pointing to cooperative effects of the different modules. We found that both the FxxA and LFDE modules of the eight BRC repeats make relevant contributions to the binding of monomeric RAD51, as evidenced by both FxxA and LFDE being associated with net disruption of binding function upon shuffling with other modules. This is also consistent with the finding by previous studies that the FxxA module in isolation (i.e. lacking the LFDE module) has only a modest affinity for RAD51^19,26,36^. Interestingly, we found that both modules often failed to act in concert to give high affinity binders within the natural repeats.

Our shuffling approach helps to pick apart the role played by both modules in their various repeats, allowing us to discern module-specific effects that are otherwise obscured when measured in their parental combinations. The BRC repeats provide a fascinating example of how Nature can exploit modularity to fine-tune function. By recycling variations on the FxxA β-hairpin and LFDE α-helix ‘themes’, a range of affinities (spanning two orders of magnitude for the natural BRC repeats) was achieved, without the need to resort to entirely novel sequences.

### Structural insight into beneficial effect of shuffling and scope for future work

The crystal structure of the RAD51:BRC8-2 complex has provided novel insight into the BRC repeat binding to RAD51, in particular in identification of the extended β-hairpin formed by the FxxA module and the critical role Ser2056^BRC8^ in facilitating the formation of this structure. The enhanced affinity of BRC8-2 may render it an attractive tool in studies that seek to investigate the effect of disrupting the RAD51:BRCA2 interaction using cell penetrating peptide derivatives of BRC repeats^36,39^. We were able to demonstrate the utility of BRC8-2 in a functional cellular assay for disruption of radiation-induced RAD51 foci formation. Strategies to further stabilize this β-hairpin may involve the use of tryptophan-tryptophan cross-strand pairs^40^ or even artificial crosslinks such as triazole^41^, to achieve an additional enhancement of binding through a reduction in conformational heterogeneity prior to complex formation (as well as improving resistance to proteolytic degradation). Our work revealed that the linker between the FxxA module and LFDE module is also likely to play a role in determining the affinity of the peptide. Our data on the importance of the linker region is further corroborated by a study showing that mutating the wildtype Val1532^BRC4^ to Ile or Phe results in respectively enhanced and diminished binding of BRC4 peptide to RAD51^42^.

In this study we exploited a shuffling library of natural repeats only, successfully obtaining affinity-enhanced variants, together with an improved understanding of the modular contributions to binding made by FxxA and LFDE. Several studies have investigated the effect of point mutations within the modules, which would be interesting to combine with our shuffling approach in future studies. These efforts are partially motivated by the therapeutic potential of blocking the BRCA2-RAD51 interaction. Nomme *et al* succeeded, through careful rational analysis and sophisticated molecular modeling techniques, in generating a BRC4 repeat peptide mutant that was 10-times more efficient in inhibiting the RAD51:ssDNA complex than the original BRC4 repeat peptide itself^26^. Similarly, Venkitaraman, Skylaris and colleagues calculated molecular mechanics energies combined with the Poisson– Boltzmann surface area continuum solvation (MM-PBSA) to successfully identify the BRC repeat binding hotspots as well as enabling an accurate prediction of relative binding free energies of the natural BRC repeats^29^. Scott *et al*. probed the contribution of individual residues in the FxxA epitope identifying changes that resulted in increased affinity towards RAD51^43^ (by up to ΔΔG = −0.64 kCal/mol). Unlike typical computational approaches, the affinity enhancement achieved by our shuffling approach requires no *a priori* knowledge of the binding mechanism. Guided by the fast evaluation of *K*_d_ values in microfluidic droplets, empirical models can be developed that yield novel insights into binding mechanism, and chimeras with improved affinity for use in various diagnostic and therapeutic applications can be obtained. Our crystal structure of a shuffled repeat in complex with monomeric RAD51 allowed detailed investigation of its binding mechanism and we would recommend that future efforts at developing BRC peptides with enhanced RAD51 affinity exploit such structural investigations.

### Conclusion

This work combines rational and combinatorial engineering productively and illustrates a general strategy: knowledge of functional units of proteins that are autonomously folded and functional, bypasses the need to design proteins from scratch, while their shuffling reduces the library complexity vastly, compared to the total sequence randomization typical in directed evolution approaches. Defining such functional modules better will provide the basis for more sophisticated libraries created by module shuffling, to ultimately reach the goal of eliciting functional proteins more quickly.

## Methods

### Reagents

Preparation of BRC4 repeat peptide, N-terminally labeled with fluorescein (BRC4^fl^, sequence CKEPTLLGFHTASGKKVKIAKESLDKVKNLFDEKEQ) was described previously^27^. CHES was from Sigma, Pico-Surf 1 was from Dolomite, HFE-7500 was from 3M.

### Plasmid constructs & cloning

For the construction of 8 parental and 56 shuffled BRC peptides, as well as several more mutant peptides, please see SI (Supplementary Figs. 1, 2 and Supplementary Table 1). The *E*. *coli* expression construct for monomeric RAD51 (pBAT4-HumRadA22), has been described previously^24^. Cloning of plasmids GFP and GFP-BRC8-2 for mammalian cell transfection is described in SI (Supplementary Fig. 3 and Supplementary Table 2). All plasmids used in this study may be requested from the corresponding authors.

### Protein expression & purification

The 64 different GB1-BRC peptide fusion constructs were separately transformed to chemically competent *E*. *coli* BL21(DE3). Overnight LB pre-cultures were used to inoculate 20 mL LB, which were grown up to mid-log phase (OD_600_ of 0.5). Expression was induced using 1 mM IPTG and cultures were incubated for a further 3 hours at 37 °C. Cells were then harvested through centrifugation and lysed by the addition of BugBuster/Benzonase lysis reagent (Novagen, with 5 mM imidazole, 20 mM Tris-HCl, 100 mM NaCl, pH 8). The resuspended mixture was incubated for 20 minutes at room temperature, then loaded directly onto a Ni-NTA protein miniprep column (His Spin Protein Miniprep, Zymo Research). Protein was washed following the manufacturer’s instructions. Proteins were eluted in 500 mM imidazole, 20 mM Tris-HCl, 100 mM NaCl, pH 8, 150 µL. Protein concentrations were quantified by UV absorption at 280 nm (using a Nanodrop spectrophotometer) and corrected for the presence of truncated side-products by SDS PAGE (Supplementary Fig. 4). Monomeric RAD51 expression and purification for fluorescence anisotropy measurements was carried out as described previously^24^, where monomeric RAD51 was called ‘HumRadA22’.

### Monomeric RAD51:BRC8-2 complex purification for crystallography

*E*. *coli* BL21(DE3) cells carrying pUBS520 plasmid for rare AGA/AGG encoding tRNA were transformed with GB1-BRC8-2 or monomeric RAD51 constructs and grown at 37 °C in 1 L of 2x YT medium in shaker flasks in the presence of 100 μg/mL of ampicillin and 25 μg/mL kanamycin until OD_600_ of 0.8. Expression was induced with 0.4 mM IPTG for three hours. Cells were resuspended in 25 mL of 50 mM Tris-HCl (pH=8.0), 100 mM NaCl, 20 mM imidazole and lysed on an Emulsiflex C5 homogenizer (Avestin). Cell lysate was centrifuged at 15 000 *g* for 30 min and supernatant collected. GB1-BRC8-2 lysate was loaded on a 3 mL Ni-NTA agarose matrix (Cube Biotech), followed by the application of monomeric RAD51 lysate. Column matrix was washed with 5 column volumes 50 mM Tris-HCl pH 8.0, 100 mM NaCl, 20 mM imidazole. Complex was eluted with 50 mM Tris-HCl pH 8.0, 100 mM NaCl, 200 mM imidazole into 2 ml fractions. Fractions containing the proteins of interest were pooled and incubated with 100 μL of 2 mg/ml TEV protease overnight at 4°C. Cleaved GB1 fusion partner was removed from the solution by a second Ni-NTA affinity step, collecting the flow-through that contains the monomeric RAD51:BRC8-2 complex. Flow-through was concentrated on a centrifugal filter (Amicon, MWCO 3000 Da) to 2 ml volume and loaded into a Superdex 75 16/60 prep grade size exclusion column (GE Lifesciences), previously equilibrated with 20 mM CHES pH 9.5, 100 mM NaCl, 1 mM EDTA. The complex eluted at 75 ml, the fractions containing the complex were pooled and the complex concentrated to 0.45 mM.

### Fluorescence polarization competition assay and microfluidic based measurements

Fluorescence polarization was measured in nanoliter droplets produced and analyzed in microfluidic devices to quantify bound vs unbound RAD51:BRC complexes and derive *K*_d_ values, essentially as described previously ^27^. Briefly, nanoliter droplets were generated from a well in which an increasing amount of peptide is allowed to compete with BRC4^fl^. A 4-channel parallelized device was used for all measurements to increase throughput. A 10x objective was used and the power of the 488 nm diode laser was 50 mW. BRC peptides were pre-loaded into PTE tubing (internal Ø, 0.38mm) to avoid cleaning syringes between runs. To this end, 40 µL of each peptide was aspirated in tubing, followed a plug of 10 µL of HFE-7500 and 0.5% Pico-Surf 1 (Sphere Fluidics). Typically, four peptides were pre-loaded in each tubing, so that 16 samples could be screened at a go. All measurements were performed in CHES buffer pH 9.5, 1% BSA with HFE-7500 oil and 0.5% Pico-Surf 1 surfactant as carrier phase. Flow rates were 3 μL/min for withdrawal and 40 μL/min for 30 seconds for peptide injection. Data was fit to a competitive binding model using 12 nM for the *K*_d_ of the monomeric RAD51:BRC4_fl_ interaction^27^.

### Crystallography of monomeric RAD51:BRC8-2 complex

Monomeric RAD51:BRC8-2 complex was crystallized using sitting-drop vapor diffusion in a 96-well MRC plate format. 40 mM ADP/Mg^2+^ water solution was added to 0.45 mM complex in a 1:9 ratio. 200 nL of the complex was then mixed with 200 nL of the crystallization condition using a Mosquito liquid handling robot (TTP Labtech). Crystals were observed in 0.2 M NH_4_Cl, 20% w/v PEG 3350 and used directly for data collection without the need for further optimization of the crystallization conditions.

A crystal was cryo-cooled in liquid nitrogen without the application of a cryo-protectant and diffraction data were collected at Diamond Light Source (Harwell, UK) synchrotron radiation source. Images were processed with autoPROC^44^. Molecular replacement phasing method was used with apo monomeric RAD51 structure (‘HumRadA22’, PDB: 5KDD) as a search model. The structure was refined without peptide first and the peptide was built into the clearly visible electron density manually (see Supplementary Fig. 10). Manual real-space refinement was done in Coot^45^ and automated refinement with phenix.refine^46^ and autoBUSTER^47^. Crystallographic data and refinement statistics are shown in Supplementary Table 5. The final model contains 7 complexes of monomeric RAD51 complexed with BRC8-2 peptide and one monomeric RAD51 chain with no peptide (chain H). Chain H has poorly defined electron density, which is likely caused by the lower number of crystal contacts it makes compared to other monomeric RAD51 molecules in the asymmetric unit. Individual atomic B-factors were not refined for chain H. The protein structure is fully defined in all of the complexes, but the peptide density had more variable quality. Chains B and J represent the best-defined monomeric RAD51:BRC8-2 complex and were used in the analysis. The coordinates and corresponding structure factors have been deposited to the PDB under accession number 6HQU. Contact area within the complex of both monomeric RAD51:BRC8-2 (6HQU) and RAD51:BRC4 (1n0w) was calculated (for atoms within 3.9 Å distance of atoms of the other binding partner) using a Pymol script written by Martin Christen (contact_surface v.3.0, available at https://pymolwiki.org/index.php/Contact_Surface). The script was adapted for Python3 using the 2to3 program (https://docs.python.org/2/library/2to3.html).

### Cell line

U2OS cell line (ATCC, HTB-96) was grown in DMEM media supplement with 10% FBS (Gibco™ Fetal Bovine Serum, 11573397) and 100 U/mL penicillin/streptomycin (15140122, Gibco) at 37 °C and 5% CO_2_.

### Transfection and cell treatment

Cells were transfected using Lipofectamine 3000 Transfection Reagent (Invitrogen) following manufacture’s protocol. Plasmid DNA and transfection reagent amounts were scaled for a 10-cm dish: 4 μg DNA and 7.75 μL Lipofectamine 3000.

Approximately 18 hours after transfection, cells were either exposed to 3 Gy caesium-137 g-irradiation (GammaCell 1000, Atomic Energy of Canada Ltd) or unirradiated and allowed to recover for 3 hours before being collected for analysis.

### Immunostaining

Coverslips were washed twice in phosphate buffered saline (PBS) before fixation with 4% paraformaldehyde/PBS for 15 minutes, then washed three times in PBS and permeabilized in 0.5% Triton-X/PBS for 7 minutes. Coverslips were washed three times in PBS, blocked for at least 30 min in 1% BSA-Fraction V (A3059-50G, Sigma-Aldrich)/PBS, and followed by 1 hr incubation at room temperature with RAD51 (RAD51 H-92, sc-8349, Santa Cruz) primary antibody diluted 1:100 in 1% BSA-Fraction V/PBS. The coverslips were washed three times with PBS, then incubated with anti-rabbit Alexa Fluor 647 secondary antibody (A21244, Invitrogen) diluted 1:500 in 1% BSA-Fraction V/PBS for 45 min in the dark at room temperature. Coverslips were wash three times in PBS, mounted on to slides using ProLong Gold Antifade Mountant with DAPI (P36941, Invitrogen) and stored at 4 °C for further analysis.

Cells were visualized using a Nikon Eclipse e-400 microscope with 60X objective. Images were processed and analyzed for GFP signal and RAD51 foci using CellProfiler 3.1.8.

### Flow cytometry

Cells were trypsinized, washed twice in PBS and fixed by gently vortexing while adding 1 mL ice-cold 70% ethanol drop-wise. Samples were stored for a minimum of 12 h at −20 °C. Prior to flow cytometry analysis, cells were spun down, washed twice in PBS and resuspended in around 0.5 mL Staining Solution (5 μg/mL propidium iodide (P3566, ThermoFisher Scientific), 100 μg/mL RNase A (R5503-100MG, Sigma-Aldrich) in PBS) and incubated for at least 30 min in the dark at room temperature. Cells were analyzed on a BD LSR II Flow Cytometer (BD Biosciences) and cell cycle profiles were generated after gating GFP positive cells using FlowJo v10.1 software. A detailed description of the gating strategy is found in Supplementary Fig. 12. Around 10,000 GFP positive cells were analysed per condition and experiment.

### Data availability

Supplementary data contains detailed descriptions of the cloning of bacterial expression constructs for the 64 shuffled BRC peptide variants, cloning of mammalian expression constructs and notes on the soluble expression of the shuffled BRC peptide variants. Also included is a description of ITC used to cross-validate the microfluidic measurements, single concentration point measurements carried out with microfluidics and exemplary titrations carried out by microfluidics. The fluorescence anisotropy data obtained for the 64 separate titrations as well as the Matlab script used in the analysis have been uploaded as separate files. The supplementary data also contains an analysis on the effect of shuffling of BRC peptides and in particular on the effect of the exact shuffle cut-off point placement. X-ray crystallography electron density maps, data collection and refinement statistics are also to be found in the supplementary data. Additional cell images highlighting the pan-nuclear signal of RAD51 are also included in the supplementary data. The coordinates and corresponding structure factors for the monomeric RAD51:BRC8-2 complex have been deposited to the PDB under accession number 6HQU.

## Acknowledgements

We thank Diamond Light Source for access to macromolecular crystallography beam line i03 (proposal 18548) and for the data that contributed to these results. We are also grateful for access to and support by the X-ray crystallographic and Biophysical Research Facility at the Department of Biochemistry, University of Cambridge. This work was supported by the BBSRC (BB/K013629/1) and the European Research Council (ERC) under the European Union’s Horizon 2020 research and innovation program (grant agreement n° 695669). LL received a Marie Curie individual fellowship from the European Commission (grant number 659029), MB a fellowship from the Schweizerischer Nationalfonds, TP a studentship from the MRC Doctoral Training Partnership and FH is an ERC Advanced Investigator (grant no. 695669). PZV and JD were supported by Cancer Research UK (C7905/A25715). CFK acknowledges funding from the UK Engineering and Physical Sciences Research Council, EPSRC (grants EP/L015889/1 and EP/H018301/1), the Wellcome Trust (grants 3-3249/Z/16/Z and 089703/Z/09/Z) and the UK Medical Research Council, MRC (grants MR/K015850/1 and MR/K02292X/1).

## Author contributions

LL, FG, MB, MH & FH conceived the study. LL, MB & TP produced recombinant proteins. FG, ER & CFK set up the fluorescence anisotropy equipment. FG carried out the high throughput fluorescence anisotropy measurements. FG & LL analyzed the fluorescence anisotropy data. TP purified the monomeric RAD51:BRC8-2 protein complex, determined its crystal structure, conducted ITC measurements and analyzed the data. MH & TP interpreted the structural data. PZ-V and JAD designed and performed the cell-based assays. LL, TP, FG, PZ-V, JAD, MH & FH wrote the paper.

## Code availability

As described previously^27^, the transformation from intensity maps into anisotropy values from image data was carried out with a custom Matlab code (https://github.com/quantitativeimaging/icetropy). The custom Matlab script used to fit a *K*_d_ for the unlabeled competitive GB1-BRC peptides can be found as a Supplementary File.

