## Supplementary Information for "Improved RAD51 binders through motif shuffling based on the modularity of BRC repeats"

#### 1. Cloning of bacterial and mammalian expression vectors

##### 1.1. Cloning of 64 (56 shuffled and 8 parental) BRC peptide expression vectors

An efficient cloning method was developed to allow rapid production of all 64 BRCA repeat recombinations fused to GB1. An oligonucleotide assembly-by-ligation procedure was designed, whereby five oligonucleotides encoded each repeat, with only one of the five extending across the two modules (Supplementary Figure 1).

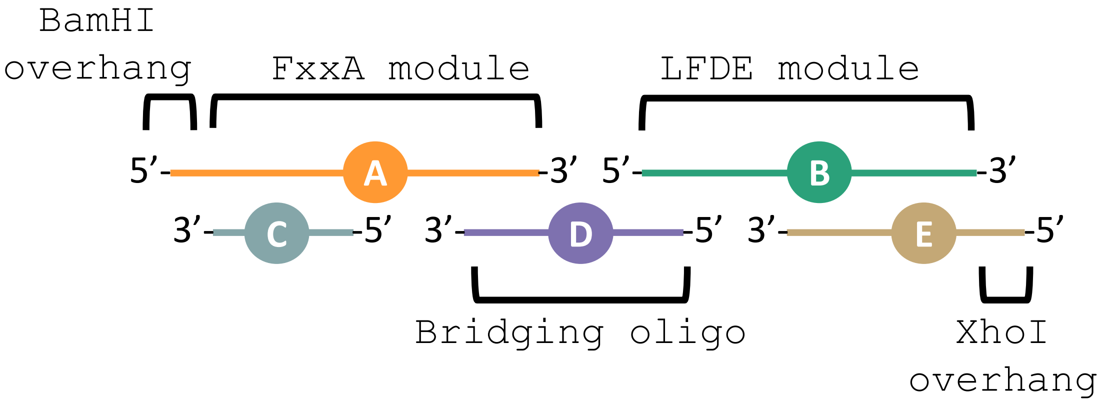

**Supplementary Figure 1**. **Schematic overview of repeat DNA library construction.** Each recombined repeat was built up from 5 oligonucleotides (A-E), designed such that only oligonucleotide D crossed the junction between modules FxxA and LFDE (bridging oligo), minimizing the total amount of synthesized material required. Oligonucleotides A and E were designed to provide 5'-single stranded vector-compatible overhangs (BamHI and XhoI, respectively).

A total of 96 oligonucleotides (Supplementary Table 1) were required for this strategy (8 each for oligonucleotide series A, B, C and E and 64 separate oligonucleotides for the D series of cross-bridging oligonucleotides). This procedure allowed repeated use of the same oligonucleotides (except the bridging oligo) in different combinations, using the following steps:

**Step 1. Phosphorylation of oligonucleotide 5’-ends**

The construction method started with the enzymatic 5’-phosphorylation of the B, C and D oligonucleotides, treated in separate reactions (30 µL for B & C, 20 µL for D). Each reaction consisted of 15 µM oligonucleotide, 0.5 units/µL T4 polynucleotide kinase (PNK, NEB) in 1x T4 ligase buffer (NEB), supplemented with an additional 30 nmol each (B & C) or 10 nmol each (D) of DTT and ATP. The reactions were incubated at 37 °C for 30 minutes, followed by heat inactivation of PNK by heating to 65 °C for 20 minutes.

**Step 2. Annealing of oligonucleotides**

Oligonucleotides A and E were mixed with the same reaction components as oligonucleotides B, C and D, at the same oligonucleotide concentration, but lacking PNK enzyme, to a volume of 30 µL. The heat-inactivated reaction mixtures for oligos B, C and D from **Step 2** were mixed with oligos A and E at 3 µM of each of the five oligonucleotides in a total volume of 17 µL. To promote annealing of the oligonucleotides, the solution was heated to 95 °C for two minutes, then incubated at 52 °C for 10 minutes, followed by cooling to 4 °C.

**Step 3. Ligation of oligonucleotides**

The 64 different series of annealed oligonucleotides from **Step 3** (i.e. A, B, C, D and E) were then supplemented with an additional 30 nmol of ATP, 30 nmol of DTT and 100 units of T4 DNA ligase (NEB), bringing the total volume of the 64 different reactions to 20 µL each. The mixtures were incubated at 16 °C for 16 hours, to allow the ligation of internal nicks (i.e. A to B, C to D and D to E) to form full-length duplexes. As the 5’-ends of oligonucleotides A and E had not been phosphorylated, formation of multimeric complexes due to self-ligation of the duplex was prevented. These duplexes were then purified by silica spin columns (DNA Clean & Concentrator, Zymo Research).

**Step 4. Insertion into circular expression vectors**

The vector pOP3BT (http://hyvonen.bioc.cam.ac.uk/pOP-vectors/) was digested with restriction enzymes BamHI and XhoI, the 5’-ends of the linearized vector were dephosphorylated using a thermosensitive alkaline phosphatase (FastAP, ThermoFisher Scientific), before the DNA was subjected to agarose gel electrophoresis, purification by silica column (DNA Clean & Concentrator, Zymo Research) and finally dialysis against MiliQ water for 30 minutes using a mixed cellulose ester membrane (MF, Millipore). To produce circular plasmids carrying the BRC repeats as C-terminal fusions to the GB1 domain, ligation reactions (5 µL) were prepared consisting of 50 ng linearized vector, 2 ng insert (equating to 5:1 molar ratio of insert:vector), 25 units T4 DNA ligase (NEB), in 1x T4 DNA ligase buffer (NEB), supplemented with 5 nmol ATP and 5 nmol DTT. The ligations were incubated at 16 °C for 16 hours, then transformed to 20 µL of chemically competent DH5alpha *E. coli* (Alpha-Select Silver Efficiency, Bioline) by heat shock protocol and plated on LB agar plates containing 50 µg/mL carbenicillin. Two colonies of each transformation were picked and grown for miniprepping and verification by Sanger sequencing. An example of a HisTag-GB1 fusion protein (for BRC4) is provided below (Supplementary Figure 2).

```
                                  atgaatggactgaatgatatctttgaagcg
                                   M  N  G  L  N  D  I  F  E  A
cagaaaattgaatggcatgaatccggatctcatcaccatcaccatcaccatcacactagt
 Q  K  I  E  W  H  E  S  G  S  H  H  H  H  H  H  H  H  T  S
acctacaaactgatcctgaacggtaaaaccctgaaaggtgaaaccaccaccgaagctgta
 T  Y  K  L  I  L  N  G  K  T  L  K  G  E  T  T  T  E  A  V
gacgctgctactgctgaaaaagttttcaaacagtacgctaacgacaacggtgtggacggt
 D  A  A  T  A  E  K  V  F  K  Q  Y  A  N  D  N  G  V  D  G
gaatggacctacgacgacgctaccaaaaccttcacggttacggaaaccggtagtggcacc
 E  W  T  Y  D  D  A  T  K  T  F  T  V  T  E  T  G  S  G  T
agtgggtcgacagaaaacctgtacttccagggatccaaaattaaagaaccaacactgctg
 S  G  S  T  E  N  L  Y  F  Q  G  S  K  I  K  E  P  T  L  L
ggctttcatactgcctccggtaagaaagtgaaaattgcaaaagaaagtctggataaagta
 G  F  H  T  A  S  G  K  K  V  K  I  A  K  E  S  L  D  K  V
aaaaacctgtttgatgaaaaagaacagggcactggctcgagctaa
 K  N  L  F  D  E  K  E  Q  G  T  G  S  S  -
```

**Supplementary Figure 2**. **Partial DNA and amino acid sequence for construct pOP3TB-BRC4.** The BamHI restriction site (GGATCC) is indicated in red, while the XhoI restriction site (CTCGAG) is indicated in blue. The amino acid sequence for GB1-BRC4 is shown below the DNA sequence. The His-tag is highlighted in yellow, the GB1 domain in green, the FxxA module of BRC4 in pink and the LFDE module of BRC4 in blue.

**Supplementary Table 1.** Sequences of oligonucleotides used in the assembly of 64 recombinant BRC peptides. All oligos were from Sigma-Aldrich (Haverhill, England) and were ordered as desalted only.

| Name oligonucleotide | Oligonucleotide sequence (5'→3') |
|---|---|
| A1_BamHI_BRC4_F | GATCCAAAATTAAAGAACCAACACTGCTGGGCTTTCATACTGCCTCCGGTAAG |
| A2_BamHI_BRC1_F | GATCCAGCAATCATAGTTTTGGTGGCAGTTTTCGCACTGCGAGTAACAAA |
| A3_BamHI_BRC2_F | GATCCGAAAATGAAGTTGGCTTTCGCGGTTTTTATAGTGCACATGGCACG |
| A4_BamHI_BRC3_F | GATCCGATTTTGAAACGTCTGATACTTTTTTTCAGACGGCATCAGGCAAA |
| A5_BamHI_BRC5_F | GATCCTCTGTAATTGAAAATTCTGCACTGGCATTTTATACCAGTTGTAGTCGC |
| A6_BamHI_BRC6_F | GATCCAATTTTGAAGTTGGCCCTCCGGCCTTTCGTATCGCGAGTGGCAAA |
| A7_BamHI_BRC7_F | GATCCTCATCCGCCAATACGTGTGGTATTTTTAGTACAGCTAGCGGTAAA |
| A8_BamHI_BRC8_F | GATCCGTAAATAGCAGCGCCTTTAGTGGCTTTAGTACCGCCTCCGGCAAA |
| B1_BRC2_XhoI_F | AAACTGAATGTGTCAACAGAAGCGCTGCAGAAAGCAGTAAAACTGTTTTCAGATATTGAAAATATTTCAGGC |
| B2_BRC6_XhoI_F | ATCGTCTGTGTGTCACATGAAACCATTAAAAAAGTTAAAGATATTTTTACCGATAGTTTTAGTAAAGTGGGC |
| B3_BRC1_XhoI_F | GAAATTAAACTGTCAGAACATAACATCAAAAAATCCAAAATGTTTTTTAAAGATATCGAAGAACAGTATGGC |
| B4_BRC3_XhoI_F | AATATTAGCGTGGCTAAAGAATCCTTTAACAAAATTGTCAATTTTTTTGATCAGAAACCAGAAGAACTGCATGGC |
| B5_BRC4_XhoI_F | AAAGTGAAAATTGCAAAAGAAAGTCTGGATAAAGTAAAAAACCTGTTTGATGAAAAAGAACAGGGCACTGGC |
| B6_BRC5_XhoI_F | AAAACATCAGTCAGTCAGACCTCACTGCTGGAAGCAAAAAAATGGCTGCGCGAAGGTATTTTTGATGGTGGC |
| B7_BRC7_XhoI_F | TCAGTTCAGGTGTCAGATGCAAGCCTGCAGAATGCCCGTCAGGTGTTTTCTGAAATCGAAGATGGC |
| B8_BRC8_XhoI_F | CAGGTGTCAATCCTGGAATCCAGTCTGCATAAAGTGAAAGGTGTGCTGGAAGAATTTGATCTGATTCGCACTGGC |
| C1_BamHI_BRC4_R | AAAGCCCAGCAGTGTTGGTTCTTTAATTTTG |
| C2_BamHI_BRC1_R | GCGAAAACTGCCACCAAAACTATGATTGCTG |
| C3_BamHI_BRC2_R | ATAAAAACCGCGAAAGCCAACTTCATTTTCG |
| C4_BamHI_BRC3_R | CTGAAAAAAAGTATCAGACGTTTCAAAATCG |
| C5_BamHI_BRC5_R | ATAAAATGCCAGTGCAGAATTTTCAATTACAGAG |
| C6_BamHI_BRC6_R | ACGAAAGGCCGGAGGGCCAACTTCAAAATTG |
| C7_BamHI_BRC7_R | ACTAAAAATACCACACGTATTGGCGGATGAG |
| C8_BamHI_BRC8_R | ACTAAAGCCACTAAAGGCGCTGCTATTTACG |
| D1_BRC4_2_R | TGACACATTCAGTTTCTTACCGGAGGCAGTATG |
| D2_BRC4_6_R | TGACACACAGACGATCTTACCGGAGGCAGTATG |
| D3_BRC1_6_R | TGACACACAGACGATTTTGTTACTCGCAGT |
| D4_BRC2_6_R | TGACACACAGACGATCGTGCCATGTGCACT |
| D5_BRC3_6_R | TGACACACAGACGATTTTGCCTGATGCCGT |
| D6_BRC5_6_R | TGACACACAGACGATGCGACTACAACTGGT |
| D7_BRC6_6_R | TGACACACAGACGATTTTGCCACTCGCGAT |
| D8_BRC7_6_R | TGACACACAGACGATTTTACCGCTAGCTGT |
| D9_BRC8_6_R | TGACACACAGACGATTTTGCCGGAGGCGGT |
| D10_BRC4_1_R | TGACAGTTTAATTTCCTTACCGGAGGCAGTATG |
| D11_BRC4_3_R | AGCCACGCTAATATTCTTACCGGAGGCAGTATG |
| D12_BRC4_4_R | TGCAATTTTCACTTTCTTACCGGAGGCAGTATG |
| D13_BRC4_5_R | ACTGACTGATGTTTTCTTACCGGAGGCAGTATG |
| D14_BRC4_7_R | TGACACCTGAACTGACTTACCGGAGGCAGTATG |
| D15_BRC4_8_R | CAGGATTGACACCTGCTTACCGGAGGCAGTATG |
| D16_BRC1_1_R | TGACAGTTTAATTTCTTTGTTACTCGCAG |
| D17_BRC1_2_R | TGACACATTCAGTTTTTTGTTACTCGCAG |
| D18_BRC1_3_R | AGCCACGCTAATATTTTTGTTACTCGCAG |
| D19_BRC1_4_R | TGCAATTTTCACTTTTTTGTTACTCGCAG |
| D20_BRC1_5_R | ACTGACTGATGTTTTTTTGTTACTCGCAG |
| D21_BRC1_7_R | TGACACCTGAACTGATTTGTTACTCGCAG |
| D22_BRC1_8_R | CAGGATTGACACCTGTTTGTTACTCGCAG |
| D23_BRC2_1_R | TGACAGTTTAATTTCCGTGCCATGTGCACT |
| D24_BRC2_2_R | TGACACATTCAGTTTCGTGCCATGTGCACT |
| D25_BRC2_3_R | AGCCACGCTAATATTCGTGCCATGTGCACT |
| D26_BRC2_4_R | TGCAATTTTCACTTTCGTGCCATGTGCACT |
| D27_BRC2_5_R | ACTGACTGATGTTTTCGTGCCATGTGCACT |
| D28_BRC2_7_R | TGACACCTGAACTGACGTGCCATGTGCACT |
| D29_BRC2_8_R | CAGGATTGACACCTGCGTGCCATGTGCACT |
| D30_BRC3_1_R | TGACAGTTTAATTTCTTTGCCTGATGCCGT |
| D31_BRC3_2_R | TGACACATTCAGTTTTTTGCCTGATGCCGT |
| D32_BRC3_3_R | AGCCACGCTAATATTTTTGCCTGATGCCGT |
| D33_BRC3_4_R | TGCAATTTTCACTTTTTTGCCTGATGCCGT |
| D34_BRC3_5_R | ACTGACTGATGTTTTTTTGCCTGATGCCGT |
| D35_BRC3_7_R | TGACACCTGAACTGATTTGCCTGATGCCGT |
| D36_BRC3_8_R | CAGGATTGACACCTGTTTGCCTGATGCCGT |
| D37_BRC5_1_R | TGACAGTTTAATTTCGCGACTACAACTGGT |
| D38_BRC5_2_R | TGACACATTCAGTTTGCGACTACAACTGGT |
| D39_BRC5_3_R | AGCCACGCTAATATTGCGACTACAACTGGT |
| D40_BRC5_4_R | TGCAATTTTCACTTTGCGACTACAACTGGT |
| D41_BRC5_5_R | ACTGACTGATGTTTTGCGACTACAACTGGT |
| D42_BRC5_7_R | TGACACCTGAACTGAGCGACTACAACTGGT |
| D43_BRC5_8_R | CAGGATTGACACCTGGCGACTACAACTGGT |
| D44_BRC6_1_R | TGACAGTTTAATTTCTTTGCCACTCGCGAT |

| | |
|---|---|
| D45_BRC6_2_R | TGACACATTCAGTTTTTTGCCACTCGCGAT |
| D46_BRC6_3_R | AGCCACGCTAATATTTTTGCCACTCGCGAT |
| D47_BRC6_4_R | TGCAATTTTCACTTTTTTGCCACTCGCGAT |
| D48_BRC6_5_R | ACTGACTGATGTTTTTTTGCCACTCGCGAT |
| D49_BRC6_7_R | TGACACCTGAACTGATTTGCCACTCGCGAT |
| D50_BRC6_8_R | CAGGATTGACACCTGTTTGCCACTCGCGAT |
| D51_BRC7_1_R | TGACAGTTTAATTTCTTTACCGCTAGCTGT |
| D52_BRC7_2_R | TGACACATTCAGTTTTTTACCGCTAGCTGT |
| D53_BRC7_3_R | AGCCACGCTAATATTTTTACCGCTAGCTGT |
| D54_BRC7_4_R | TGCAATTTTCACTTTTTTACCGCTAGCTGT |
| D55_BRC7_5_R | ACTGACTGATGTTTTTTTACCGCTAGCTGT |
| D56_BRC7_7_R | TGACACCTGAACTGATTTACCGCTAGCTGT |
| D57_BRC7_8_R | CAGGATTGACACCTGTTTACCGCTAGCTGT |
| D58_BRC8_1_R | TGACAGTTTAATTTCTTTGCCGGAGGCGGT |
| D59_BRC8_2_R | TGACACATTCAGTTTTTTGCCGGAGGCGGT |
| D60_BRC8_3_R | AGCCACGCTAATATTTTTGCCGGAGGCGGT |
| D61_BRC8_4_R | TGCAATTTTCACTTTTTTGCCGGAGGCGGT |
| D62_BRC8_5_R | ACTGACTGATGTTTTTTTGCCGGAGGCGGT |
| D63_BRC8_7_R | TGACACCTGAACTGATTTGCCGGAGGCGGT |
| D64_BRC8_8_R | CAGGATTGACACCTGTTTGCCGGAGGCGGT |
| E1_BRC6_XhoI_R | TCGAGCCCACTTTACTAAAACTATCGGTAAAAATATCTTTAACTTTTTTAATGGTTTCATG |
| E2_BRC2_XhoI_R | TCGAGCCTGAAATATTTTCAATATCTGAAAACAGTTTTACTGCTTTCTGCAGCGCTTCTGT |
| E3_BRC1_XhoI_R | TCGAGCCATACTGTTCTTCGATATCTTTAAAAAACATTTGGATTTTTTGATGTTATGTTC |
| E4_BRC3_XhoI_R | TCGAGCCATGCAGTTCTTCTGGTTTCTGATCAAAAAAATTGACAATTTTGTTAAAGGATTCTTT |
| E5_BRC4_XhoI_R | TCGAGCCAGTGCCCTGTTCTTTTTCATCAAACAGGTTTTTACTTTATCCAGACTTTCTTT |
| E6_BRC5_XhoI_R | TCGAGCCACCATCAAAAATACCTTCGCGCAGCCATTTTTTTGCTTCCAGCAGTGAGGTCTG |
| E7_BRC7_XhoI_R | TCGAGCCATCTTCGATTTCAGAAAACACCTGACGGGCATTCTGCAGGCTTGCATC |
| E8_BRC8_XhoI_R | TCGAGCCAGTGCGAATCAGATCAAATTCTTCCAGCACACCTTTCACTTTATGCAGACTGGATTC |

##### 1.2. Cloning of further bacterial plasmid expression constructs

The construct pOP3BT-BRC4 del K1530 was produced as pOP3BT-BRC4 described above, except that instead of oligonucleotides A1_BamHI_BRC4_F and D12_BRC4_4_R, oligonucleotides A_BamHI_BRC4del_F (5'-GATCCAAAATTAAAGAACCAACACTGCTGGGCTTTCATACTGCCTCCGGT) and D_BRC4_4del_R (5'-CATACTGCCTCCGGTCAGGTGTCAATCCTG) were used. To generate pOP3BT-BRC8-2$^{S2056A}$, a PCR reaction was carried out using NEBNext Ultra II Q5 Master Mix (NEB) with 60 ng pOP3BT-BRC8-2 plasmid template and 50 pmol each of mutagenic oligonucleotides in a total volume of 50 µL. The mutagenic oligonucleotides were 8_2-1_F (5'-CAGCGCCTTTGCCGGCTTTAGTACCGCCTCCGGC) and 8_2-1_R (5'-CGGCAAAGGCGCTGCTATTTACGGATCC). The thermal cycling conditions were an initial 30 second step at 98 °C, followed by 12 cycles between 10 seconds at 98 °C and 6 minutes at 65 °C, followed by a final step at 65 °C for 10 minutes, after which the reaction was held at 4 °C. To restrict methylated plasmid template, 2 µL of FastDigest DpnI (Fermentas) was added to the unpurified PCR reaction, which was incubated at 37 °C for one hour. DNA from the reaction was purified by silica column (see above). The eluate was transformed to DH5alpha *E. coli* (see above) and plated on LB agar plates containing 50 µg/mL carbenicillin. Two colonies were picked and grown for miniprepping and verification by Sanger sequencing.

##### 1.3. Cloning of mammalian expression constructs

The GFP construct (with NLS only) was prepared from the pEGFP-C1 mammalian expression vector by appending a SV40 nuclear localization signal (PKKKRKV) at the 5' of the GFP sequence, followed by a modified multiple cloning site for subsequent cloning of BRC peptide sequences. The insert was PCR-assembled using oligonucleotides EGFP_001 and EGFP_002 (Supplementary Table 2) and cloned into BglII and BamHI-digested pEGFP-C1 using sequence and ligation independent cloning (SLIC). GFP-BRC8-2 was then prepared by digesting the GFP construct with BamHI and inserting the peptide-coding sequence which was amplified from the corresponding bacterial expression construct pOP3BT-BRC8-2 using oligonucleotides EGFP_82_001 and EGFP_82_002 (Supplementary Table 2).

**Supplementary Table 2**. Oligonucleotides used for cloning of GFP only and GFP-BRC8-2 mammalian expression constructs

| .Name oligonucleotide | Oligonucleotide sequence (5'→3') |
|---|---|
| EGFP_001 | TACAAGTCCGGACTCAGATCTCCGAAGAAGAAGAGGAAGGTGGGCGGATCCTAATAACCTAG |
| EGFP_002 | TTATCTAGATCCGGTGGATCAAGCTTAGCTCGAGCCTAGGTTATTAGGATCCGCCCACC |
| EGFP_82_001 | AGAGGAAGGTGGGCGGATCCGTAAATAGCAGCGCCTTT |
| EGFP_82_002 | AGCCTAGGTTATTAGGATCCTGAAATATTTTCAATATCTGAAAACA |

**a**

```
CGCGATCACATGGTCCTGCTGGAGTTCGTGACCGCCGCCGGGATC
 R  D  H  M  V  L  L  E  F  V  T  A  A  G  I
ACTCTCGGCATGGACGAGCTGTACAAGTCCGGACTCAGATCTCCG
 T  L  G  M  D  E  L  Y  K  S  G  L  R  S  P
AAGAAGAAGAGGAAGGTGGGCGGATCCTAATAACCTAGGCTCGAG
 K  K  K  R  K  V  G  G  S  *  -  -  -  -  -
CTAAGCT
 -  -
```

**b**

```
CGCGATCACATGGTCCTGCTGGAGTTCGTGACCGCCGCCGGGATC
 R  D  H  M  V  L  L  E  F  V  T  A  A  G  I
ACTCTCGGCATGGACGAGCTGTACAAGTCCGGACTCAGATCTCCG
 T  L  G  M  D  E  L  Y  K  S  G  L  R  S  P
AAGAAGAAGAGGAAGGTGGGCGGATCCGTAAATAGCAGCGCCTTT
 K  K  K  R  K  V  G  G  S  V  N  S  S  A  F
AGTGGCTTTAGTACCGCCTCCGGCAAAAAACTGAATGTGTCAACA
 S  G  F  S  T  A  S  G  K  K  L  N  V  S  T
GAAGCGCTGCAGAAAGCAGTAAAACTGTTTTCAGATATTGAAAAT
 E  A  L  Q  K  A  V  K  L  F  S  D  I  E  N
ATTTCAGGATCCTAATAACCTAGGCTCGAGCTAAGCT
 I  S  G  S  *  -  -  -  -  -  -  -
```

**Supplementary Figure 3. Partial DNA and amino acid sequences for the mammalian expression constructs. a** GFP construct showing the C-terminus of GFP (green), the nuclear localization signal (turquoise) and BamHI restriction site (red). **b** GFP-BRC8-2 construct with coloring as in **a**, with the inserted FxxA module of BRC8 highlighted in pink and the LFDE module of BRC2 highlighted in blue.

#### 2. Notes on soluble expression of BRC peptides

Initial attempts at studying recombinations of the BRC peptides were hindered by the poor expression of some of the non-native recombinations in *E. coli*. Although maltose binding protein (MBP)-BRC4 fusion expressed well and could be purified at high yield in *E. coli* [1], it was found that expression of MBP-BRC4-6 and MBP-BRC4-8 led to dramatically lower yields. In fact, *E. coli* BL21(DE3) transformed with MBP-BRC4-8 failed to grow at all, indicating toxicity of the expressed proteins (results not shown). As an alternative to the MBP tag, the B1 domain of Protein G (GB1, 12 kDa) has been shown to be especially efficient at enhancing the soluble expression of small proteins [2]. Indeed, expression of an initial set of 8 chimeric fusion constructs of BRC peptide to GB1 yielded 1.5-21 mg/litre, corresponding to at least 150 µL of 20 µM protein-fusion from 20 mL of bacterial culture. Densitometric analysis of SDS-PAGE was used to account for the percentage of full-length protein in each case (Supplementary Figure 4).

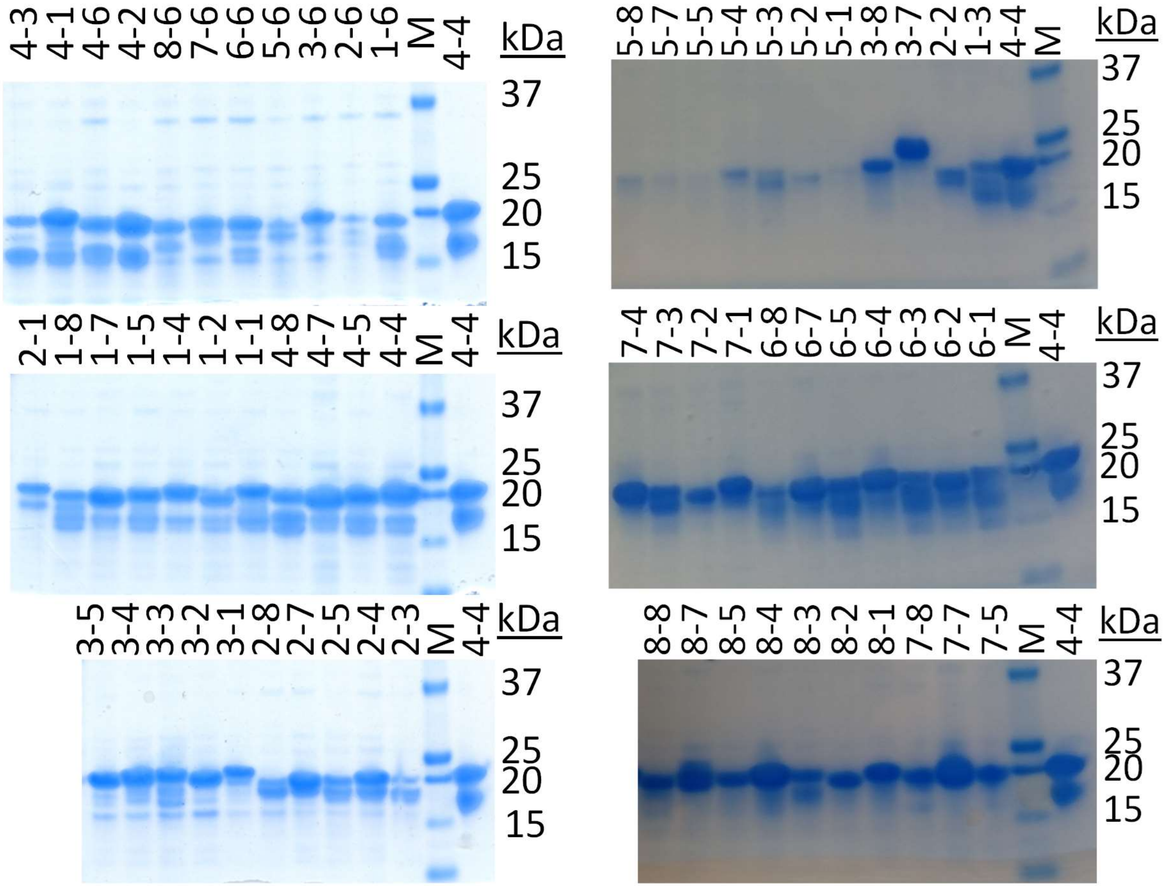

**Supplementary Figure 4. SDS-PAGE analysis of all 64 BRCA2 peptide fusions to GB1 domain variants.** Bis-Tris gels (4-12% polyacrylamide gradient) were used with a MES running buffer. The "M" denotes a protein standard ladder, of which band sizes are indicated to right of each gel on kDa. A densitometric analysis was used to correct the UV-Vis absorbance-determined concentrations for fraction of truncated protein.

#### 3. Cross-validation of anisotropy assay by isothermal titration calorimetry (ITC)

##### 3.1. ITC measurements

To determine whether the fluorescence anisotropy assay provides accurate values for BRC repeat affinities and ranks them correctly relative to each other, we selected four peptides – BRC2, BRC4, BRC8 and BRC8-2 – and measured their binding constants for monomeric RAD51 using ITC. For this purpose, we purified cleaved versions of each peptide, lacking the GB1 fusion protein. This allowed separation of any protease degradation by-products by reverse phase chromatography (RPC). Each of the peptides eluted as a single peak on a C18 RPC column and their correct molecular weight was confirmed by LCMS. To investigate whether there was any effect of the GB1 fusion on binding, a measurement using intact GB1-BRC4 was also performed. The ITC data and the fitted $K_d$ values are presented in Supplementary Figure 5a-f. BRC2 peptide was found to have a $K_d$ of 14.5 nM by ITC, which is approximately 20 times smaller than the $K_d$ measured with the microfluidic assay. A ten-fold difference was observed for BRC8 ($K_d$ of 56.5 nM compared to 573 nM), while BRC4 was found to display affinity that was four-fold higher when measured by ITC ($K_d$ of 9.3 nM vs 38 nM). The presence of the GB1 fusion did not affect binding to monomeric RAD51, as BRC4 alone and GB1-BRC4 had very similar affinities. We were unable to accurately measure the $K_d$ of the shuffled BRC8-2 peptide, as it seemed to have sub-nanomolar affinity for monomeric RAD51, which resulted in a very steep midpoint transition. Overall, it was evident that ITC ranked the affinities of BRC repeats in a similar order to fluorescence anisotropy measurements but resulted in $K_d$ values that were an order of magnitude smaller. Partial loss of peptides in the carrier oil phase used for the microfluidic screen, and likely similar in magnitude across all peptides, may partially explain this discrepancy.

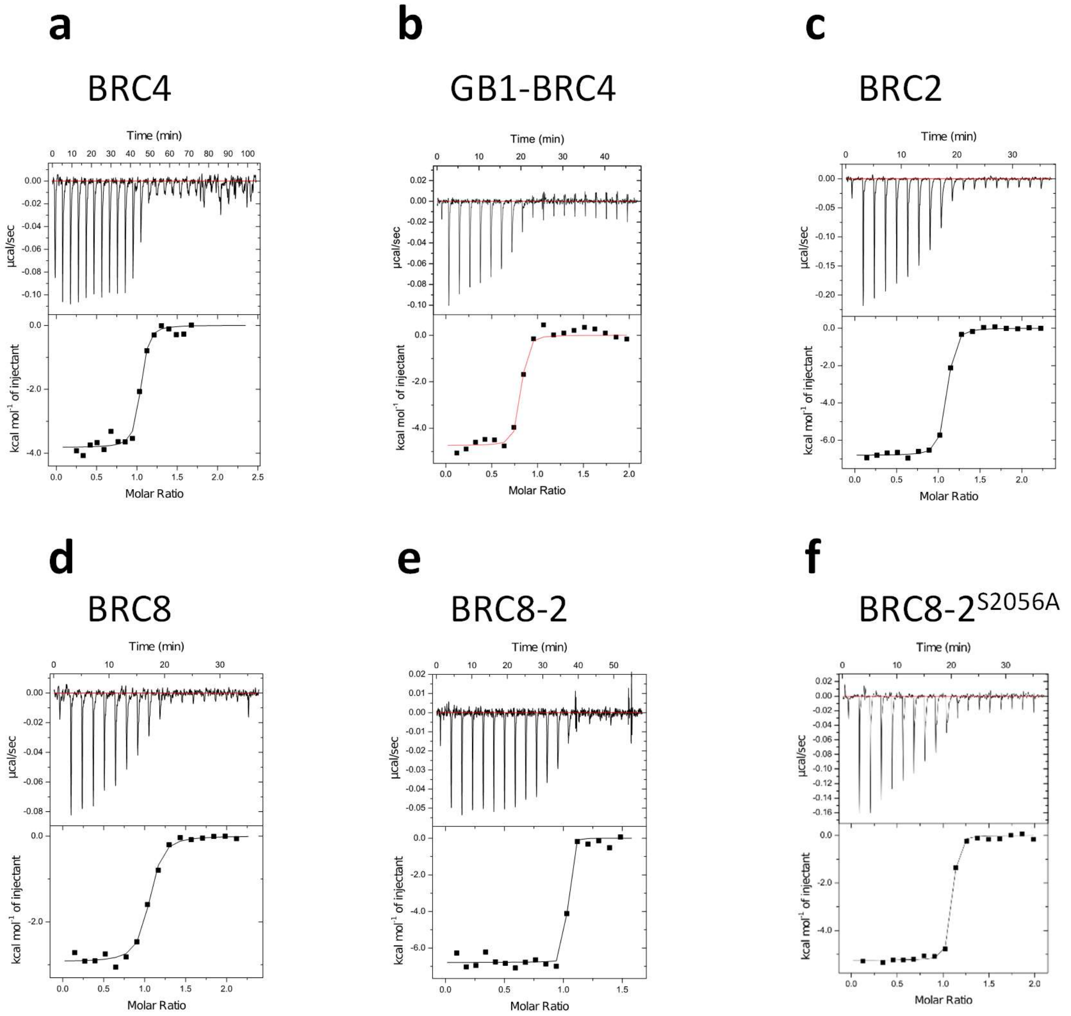

g

| | BRC4 | GB1-BRC4 | BRC2 | BRC8 | BRC8-2 | BRC8-2$^{S2056A}$ |
|---|---|---|---|---|---|---|
| Fluorescence anisotropy $K_d$, nM | 38 | 38 | 278 | 573 | 6 | - |
| $K_d$, nM | 9.3 | 11.4 | 14.5 | 56.5 | < 1 nM[a] | 5.0 |
| ΔH,, kcal/mol | -3.82 | -4.74 | -6.37 | -2.93 | -6.79 | -5.25 |
| ΔS, cal/mol/K | 23.9 | 20.4 | 15.8 | 23.3 | - | 20.4 |
| [monomeric RAD51] in cell, µM | 5 | 10 | 10 | 10 | 5 | 10 |
| [peptide] in syringe, µM | 50 | 100 | 100 | 100 | 35 | 100 |

[a] The high c-value ([RAD51 in cell]/$K_d$) of >5000, prevented an accurate determination of affinity.

**Supplementary Figure 5. ITC measurements of BRC peptide binding to monomeric RAD51. a-f** ITC data for different BRC peptides binding to monomeric RAD51. Top panel in each graph shows the baseline-corrected titration while the bottom panel shows the integrated heats of binding, together with a fit assuming 1:1 stoichiometry. **g** Table summarizing ITC measurements and observed $K_d$. Concentrations of monomeric RAD51 and peptide used in each measurement are reported to aid in the interpretation of titration data.

##### 3.2. Purification of BRC repeats with removed GB1 fusion tag

Cells carrying GB1-BRC expression constructs were grown and lysate prepared as reported in *Materials and Methods: Monomeric RAD51:BRC8-2 complex purification for crystallography*. Filtered lysate was loaded on a gravity column containing 3 mL Ni-NTA agarose matrix (Cube Biotech). Column matrix was washed with 5 column volumes 50 mM Tris-HCl pH 8.0, 100 mM NaCl, 20 mM imidazole. GB1-BRC fusion proteins were eluted with 8 mL of 50 mM Tris-HCl pH 8.0, 100 mM NaCl, 200 mM imidazole and incubated with 100 µL of 2 mg/ml TEV protease overnight at 4°C. Cleaved GB1 fusion partner was removed from the solution by a second Ni-NTA affinity step, collecting the flow-through that contains the cleaved peptide. Samples were acidified by addition of 1/10$^{th}$ volume of 100% MeCN, 1% TFA. The pH was further adjusted to pH=2 by addition of concentrated HCl. Sample was loaded on an ACE C8-300 250x4.6 mm RPC column (Hichrom). Weakly bound contaminants were washed off from the column with 4 CV of 10% MeCN, 0.1% TFA. Peptides were eluted with a 20 CV 0-100 % gradient of 90% MeCN, 0.1% TFA. Fractions containing the peptide were pooled together and diluted 1:1 by volume to halve the concentration of MeCN. Sample was loaded onto C18 RPC column (Vydac) and purified using the same conditions. Fractions containing the purified peptide were pooled and lyophilized. Peptide purity and molecular weight were confirmed by LCMS. Supplementary Figure 6 shows a represented C18 RPC chromatogram and LCMS run for BRC8-2.

**a** **b**

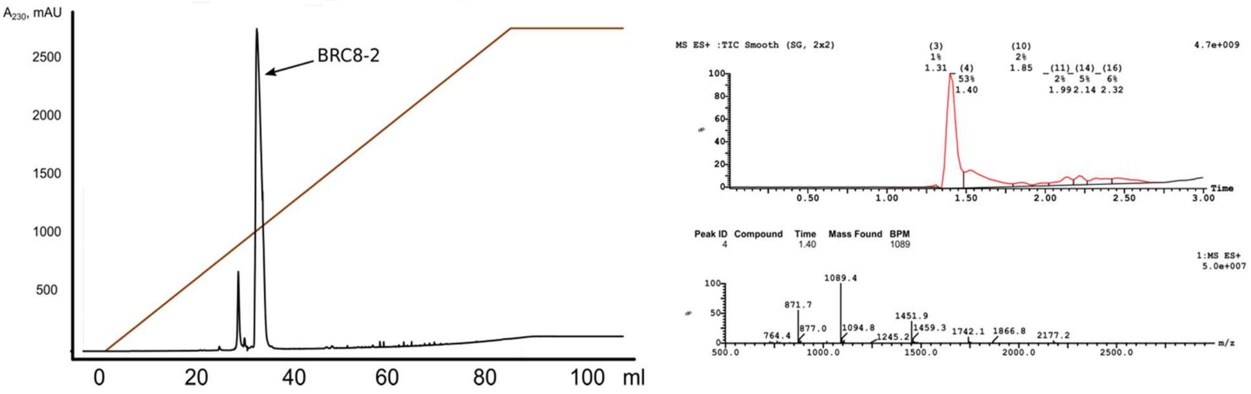

**Supplementary Figure 6. Purification of untagged BRC peptides. a** Reverse-phase chromatography of BRC8-2 on a C18 column. Major peak containing the peptide was collected and lyophilized. **b** LCMS analysis of purified BRC8-2 confirms the correct $M_W$ of the BRC8-2 peptide (4351.79 Da, including N-terminal GS- and C-terminal -GSS linkers).

##### 3.3. ITC methods

Measurements were performed with a Microcal ITC200 instrument (GE Healthcare) for peptides BRC2, BRC8, and BRC8-2 and a Microcal VP-ITC (Malvern) for BRC4. Monomeric RAD51 was buffer-exchanged into CHES pH 9.5, 100 mM NaCl, 1 mM EDTA using an Amicon spin concentrator with a 10 kDa MWCO. The flow through from monomeric RAD51 buffer exchange was used to dilute peptides resuspended in MiliQ water and water was used to match the buffer composition of the protein to minimize heats of dilution. Peptide concentrations were measured on a Nanodrop One UV/Vis spectrophotometer (at λ=205 nm) using the Scopes method. Protein and peptide concentrations were optimised in such a way as to decrease the c-value while not losing too much signal. Integration of thermogram peaks and fitting of the data was done using the ITC package in Origin 7.0 (Originlab). When fitting the binding isotherm, peptide concentration was adjusted to match a stoichiometry of N=1. Baseline instability was commonly observed after saturation during later injections, this is possibly due to peptide aggregation. Data points affected by baseline spikes were omitted from the analysis.

### 4. Microfluidic measurements of BRCA2-monomeric RAD51 interactions

#### 4.1. Single concentration point measurements to determine titration range

**Supplementary Table 3. Single concentration point measurements obtained by microfluidics-based fluorescence anisotropy competition assay.** This was carried out to ensure each chimera was measured at an optimal concentration range giving rise to the most reliable binding data in the subsequent assay. This initial measurement at 100-fold dilution of Ni-NTA purified peptides (and 100 nM BRC4$^{fl}$ with 150 nM monomeric RAD51) allowed a rough classification between strong, intermediate and weak binders. Values represent the percentage of BRC4$^{fl}$-peptide bound to monomeric RAD51, normalized to a control sample lacking competitor (100% BRC4$^{fl}$-peptide bound to monomeric RAD51) and a sample lacking monomeric RAD51 (0% BRC4$^{fl}$-peptide bound).

| BRC | 1 | 2 | 3 | 4 | 5 | 6 | 7 | 8 |
|---|---|---|---|---|---|---|---|---|
| 1 | 64 | 18 | 58 | 89 | 3 | 29 | 53 | 94 |
| 2 | 58 | 85 | 100 | 97 | 9 | 59 | 80 | 100 |
| 3 | 19 | 3 | 56 | 10 | 3 | 13 | 14 | 33 |
| 4 | 54 | 46 | 97 | 94 | 5 | 48 | 48 | 93 |
| 5 | 9 | 6 | 47 | 30 | 3 | 18 | 3 | 26 |
| 6 | 0 | 6 | 20 | 7 | 8 | 0 | 31 | 3 |
| 7 | 18 | 11 | 39 | 75 | 9 | 24 | 23 | 53 |
| 8 | 9 | 0 | 21 | 18 | 3 | 13 | 7 | 20 |

#### 4.2. Full titrations of 16 exemplary BRC repeats shown in individual graphs

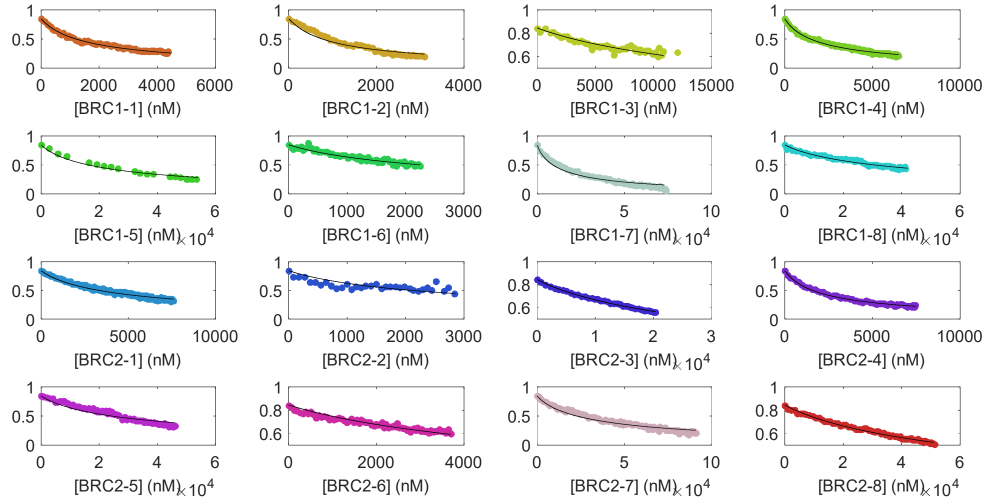

**Supplementary Figure 7. Full titrations of 16 exemplary individual GB1-BRC repeat chimeras against monomeric RAD51 / BRC4$^{fl}$-peptide measured using microfluidic setup**. This is the same data as shown in Figure 2a, but plotted as single graphs per variant, to allow closer inspection of data. As in Figure 2a, the Y-axis represents fraction BRC4$^{fl}$-peptide bound to monomeric RAD51.

#### 5. Analysis of the effect of shuffling modules from different BRCA2 repeats

##### 5.1. Gibbs free energy (ΔG) analysis of binding

To calculate Gibbs free energy, we used equation 1,

$$\Delta G = RT\ln K_d \tag{1}$$

, where R is the gas constant (1.987×$10^{-3}$ kcal/K/mol), T the temperature in Kelvin (293.15) and $K_d$ the dissociation constant measured for all 64 BRC recombinant peptide binding to monomeric RAD51 (Supplementary Table 4). To calculate $\Delta\Delta G_{parental}$ (shown in Figure 2c), equation 2 was employed:

$$\Delta\Delta G_{parental} = \Delta G_{variant} - \frac{\Delta G_{FxxA\ parent} + \Delta G_{LFDE\ parent}}{2} \tag{2}$$

, where $\Delta G_{variant}$ was the value as calculated by equation 1 for any of the 56 shuffled variants and $\Delta G_{FxxA\ parent}$ and $\Delta G_{LFDE\ parent}$ were the values as calculated by equation 1 for the natural repeats corresponding to the FxxA and LFDE 'parent' of that particular variant. The net-disruption/enhancement of binding, resulting from the shuffling in of a particular FxxA or LFDE module in combination with all other LFDE or FxxA modules, respectively, was also calculated as the row-wise (for FxxA) or column-wise (for LFDE) average of $\Delta\Delta G_{parental}$ values in Figure 2c. This value was named as $\Delta\Delta G_{module\text{-}type\ \&\ number}$ (e.g. $\Delta\Delta G_{FxxA3}$ for the FxxA module of BRC3).

**Supplementary Table 4. Thermodynamic analysis of the binding of the 64 different BRC repeats to monomeric RAD51.** ΔG values, in kCal/mol, were calculated using Equation 1 from the $K_d$ values provided in Figure 2b in the main article. The values for the 8 natural repeats are indicated in bold.

| <---LFDE modules---> \ <---FxxA modules---> | 1 | 2 | 3 | 4 | 5 | 6 | 7 | 8 |
|---|---|---|---|---|---|---|---|---|
| 1 | **-9.15** | -8.52 | -9.36 | -10.05 | -7.21 | -8.18 | -8.80 | -10.36 |
| 2 | -9.44 | **-8.79** | -10.73 | -10.58 | -7.39 | -9.06 | -10.00 | -11.03 |
| 3 | -7.44 | -7.23 | **-8.87** | -8.11 | -6.79 | -6.74 | -7.41 | -9.20 |
| 4 | -9.02 | -8.97 | -10.49 | **-9.95** | -7.34 | -7.81 | -8.83 | -10.42 |
| 5 | -7.59 | -7.35 | -8.65 | -7.62 | **-7.76** | -7.46 | -7.27 | -8.86 |
| 6 | -8.75 | -8.13 | -7.70 | -7.84 | -6.81 | **-6.72** | -8.52 | -7.37 |
| 7 | -7.93 | -7.39 | -8.38 | -8.96 | -7.65 | -7.64 | **-7.97** | -8.41 |
| 8 | -7.21 | -6.84 | -7.99 | -8.13 | -7.48 | -7.54 | -6.64 | **-8.37** |

##### 5.2. SCHEMA analysis

Using the PDB file 1n0w, together with an alignment of all eight natural BRC repeats, a residue contact map was generated (Supplementary Figure 8) using the schemacontacts.py program available on the Arnold group website (https://cheme.che.caltech.edu/groups/fha/Software.htm). Next the RASPP (Recombination As a Shortest-Path Problem) program was applied, with the minimal fragment length set to 10 residues and this script suggested the optimal crossover point to be between positions Ile1534$^{BRC4}$ and Ala1535$^{BRC4}$ (or equivalent in other repeats), with a relatively low <E> value of 0.3438, implying more functional chimeras could be produced using this cross-over (Supplementary Figure 8).

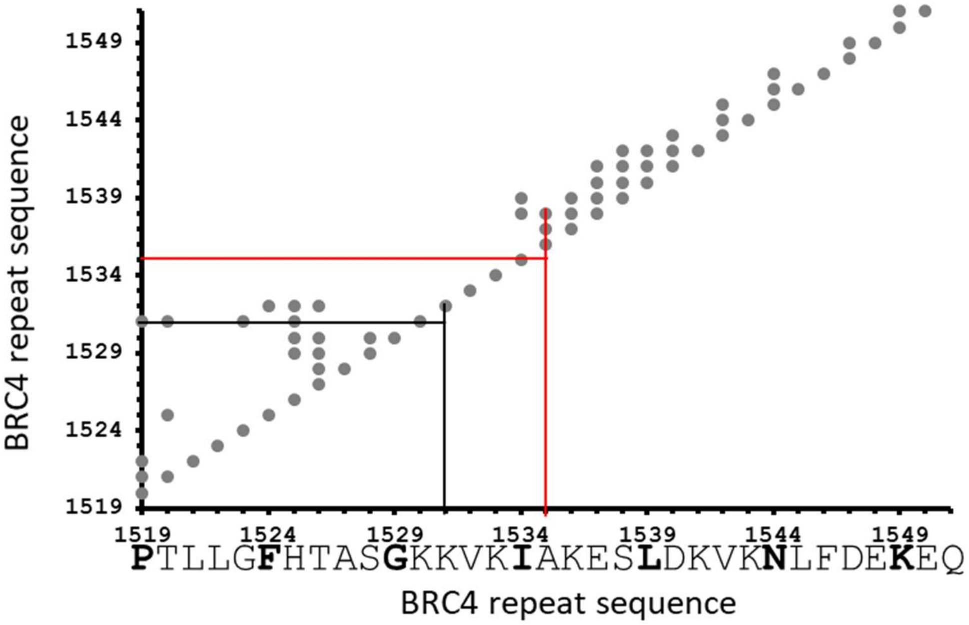

**Supplementary Figure 8. BRC peptide contact map generated by the SCHEMA algorithm**[3]. The numbering on both axes is based on the numbering of the BRC4 repeat within BRCA2. The intersection of the black lines represents the crossover point chosen in the present study, while the red lines represent the crossover point determined as being optimal by the SCHEMA algorithm.

##### 5.3. Effect of placement of shuffle cut-off point between modules FxxA and LFDE

The affinity of the interaction between peptides BRC4 and BRC4 del K1530 and monomeric RAD51 (Supplementary Figure 9a) were measured by plate reader (Pherastar, BMG). Both peptides were purified by RP-HPLC as described in section 3.2 above. The BRC4 peptide was prepared from an MBP fusion [1] and did not have the GSS C-terminal amino acids, while the BRC4 del K1530 peptide was prepared from a GB1 fusion. Correct mass was confirmed by MALDI analysis (Supplementary Figure 9b) and peptide concentration was determined by amino acid analysis (not shown). The binding competition-type assay was performed using 10 nM fl-BRC4 peptide, 15 nM monomeric RAD51 protein in 20 mM CHES (pH 9.5), 100 mM NaCl, 1 mM EDTA and 10 mg/mL BSA, with 384-well plates (black, low volume, low binding, Corning), with a total assay volume of 25 µL. Serial two-fold dilutions of GB1-BRC peptide fusions were carried out in a 96-well plate. The excitation and emission filters were 485 and 520 nm respectively. Data analysis was identical to the method used for the microfluidic measurements (Supplementary Figure 9c). Fitting to the competitive binding model gave a $K_d$ of 21 nM for BRC4 peptide and 4.1 µM for BRC4 del K1530.

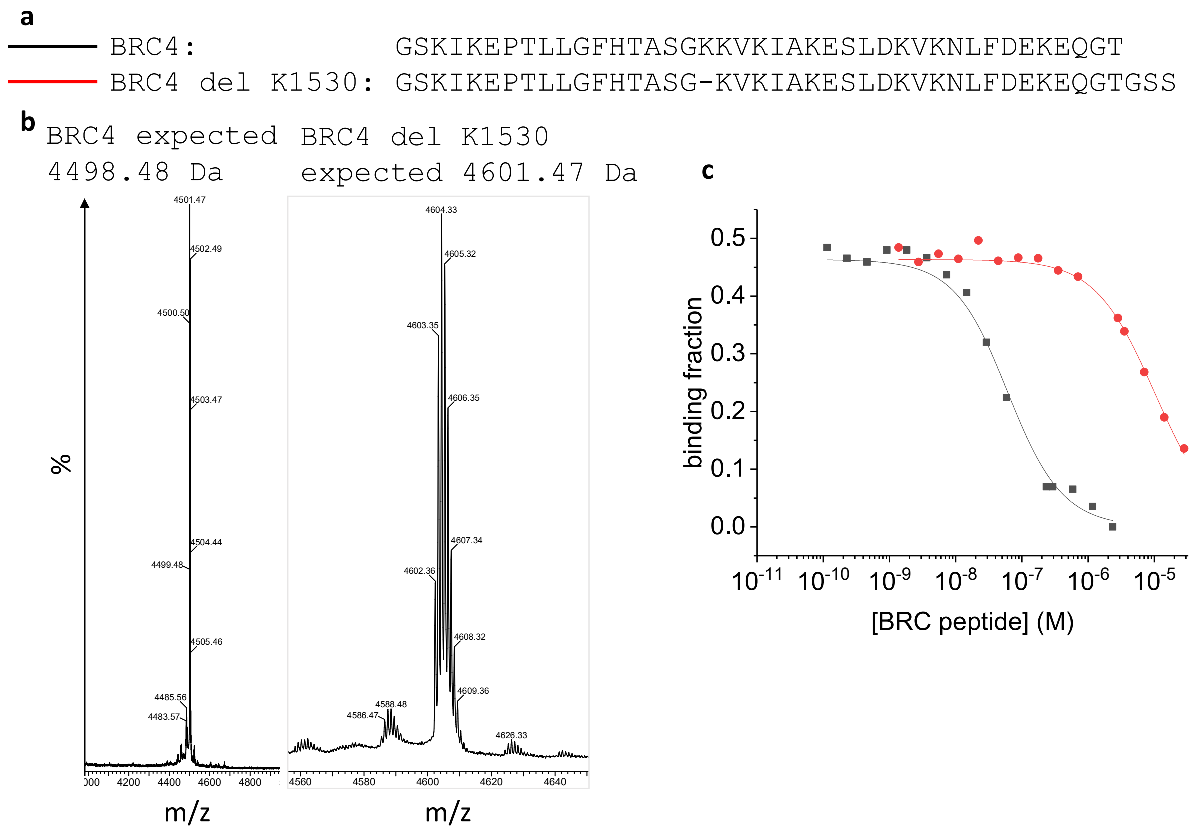

**Supplementary Figure 9. Fluorescence anisotropy competition assay to assess the effect of deletion of a residue at the shuffle cut-off point. a** Sequence alignment of BRC4 peptides with and without deletion of a central lysine (Lys1530$^{BRC4}$). **b** MALDI mass spectra confirming correct masses for both peptides after RP-HPLC purification. **c** Fluorescence anisotropy competition assay carried out in a Pherastar plate reader (BMG Labtech), with BRC4 (black) and BRC4 del K1530 (red).

#### 6. X-ray crystallography of monomeric RAD51:BRC8-2 complex

**Supplementary Table 5. X-ray crystallographic data collection and refinement statistics**

| 6HQU | |
|---|---|
| **Data processing** | |
| Wavelength (Å) | 0.97622 |
| Space group | P 1 $2_1$ 1 |
| Data collection temperature (K) | 100 |
| a, b, c (Å) | 114.403 75.473 114.792 |
| α, β, γ (°) | 90.00 97.06 90.00 |
| Resolution range (high resolution bin) (Å) | 85.870 - 1.966 (1.972-1.966) |
| $R_{merge}$ | 0.070 (1.971) |
| $R_{meas}$ | 0.091 (2.778) |
| Completeness (%) | 99.8 (100.0) |
| Number of total / unique reflections | 522829 / 137994 |
| Redundancy | 3.8 (3.9) |
| $\langle I/\sigma(I)\rangle$ | 7.4 (0.5) |
| $CC_{1/2}$ | 0.998 (0.272) |
| **Refinement** | |
| $R_{cryst}/R_{free}$ | 0.263/ 0.271 |
| Resolution range (Å) | 85.870 - 1.966 |
| Number of reflections: work/test set | 137876/ 6853 |
| Number of protein atoms | 13996 |
| Number of other atoms | 438 |
| Mean/Wilson B-factor | 68.12/ 45.47 |
| Ramachandran favoured/allowed/outliers | 1699 (99.0%) / 18 (1.0%) / 0 (0.0%) |
| RMSD bonds (Å) | 0.011 |
| RMSD angles (°) | 1.283 |

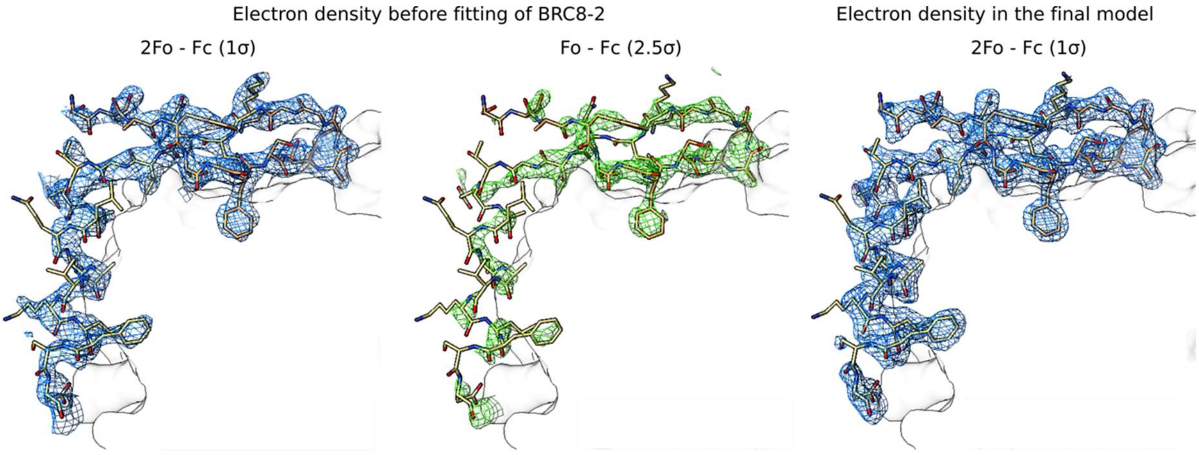

**Supplementary Figure 10. Electron density maps of bound BRC8-2 peptide.** Weighted 2Fo-Fc and Fo-Fc maps shown in blue (contoured at 1σ) and green (2.5 σ) in the left and middle columns, respectively, for BRC8-2 in chain J before it was modelled into the structure. The right column shows the final weighted 2Fo-Fc map for chain J. The electron density is clearly defined for most of the peptide, with unresolved flexible termini presumably not involved in interface formation.

#### 7. Additional data on RAD51 foci formation

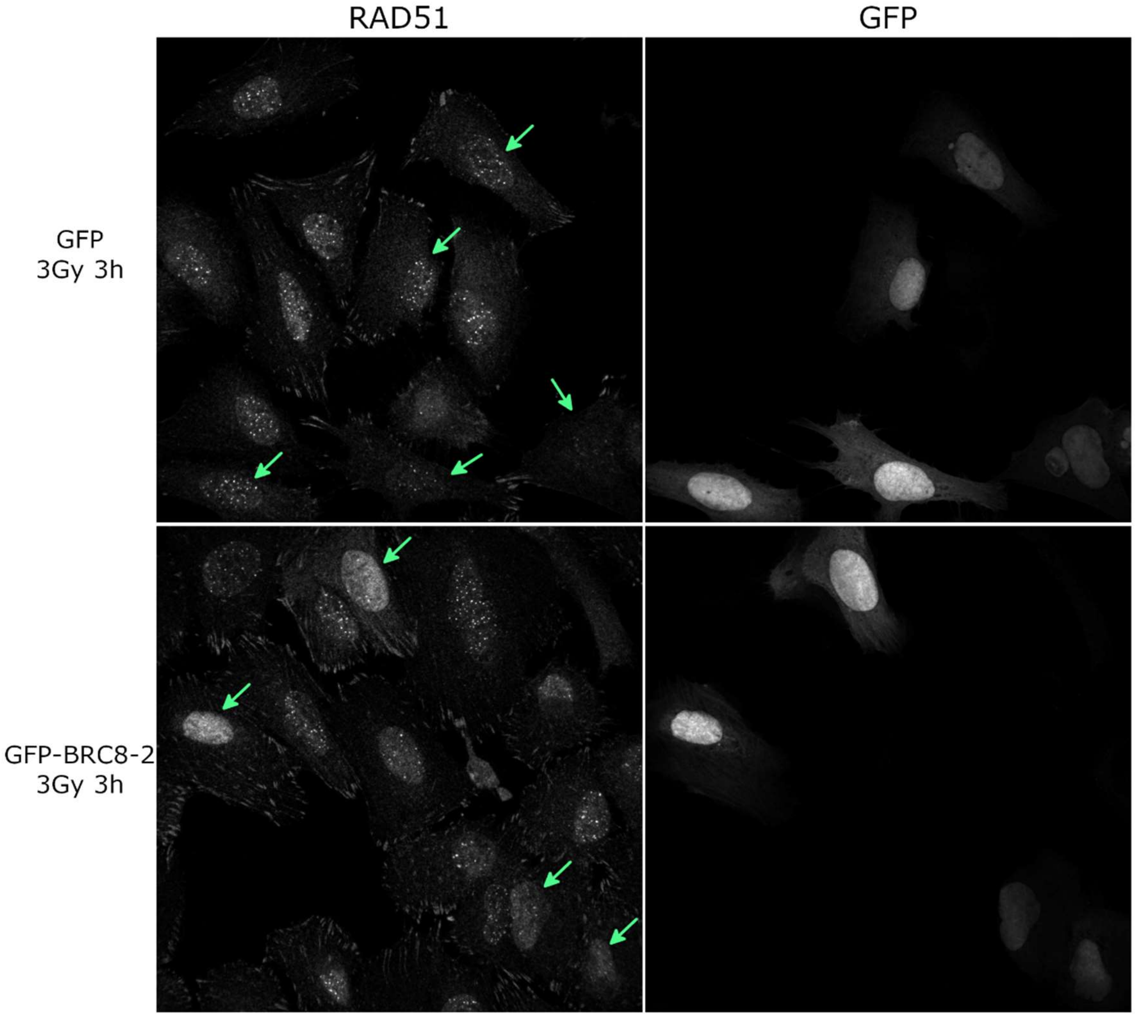

**Supplementary Figure 11. The pan-nuclear signal of RAD51 in GFP-BRC8-2 expressing U2OS cells is greater than in the GFP control cells.** Labels and conditions are the same as for Figure 4a. The arrows indicate GFP-positive cells.

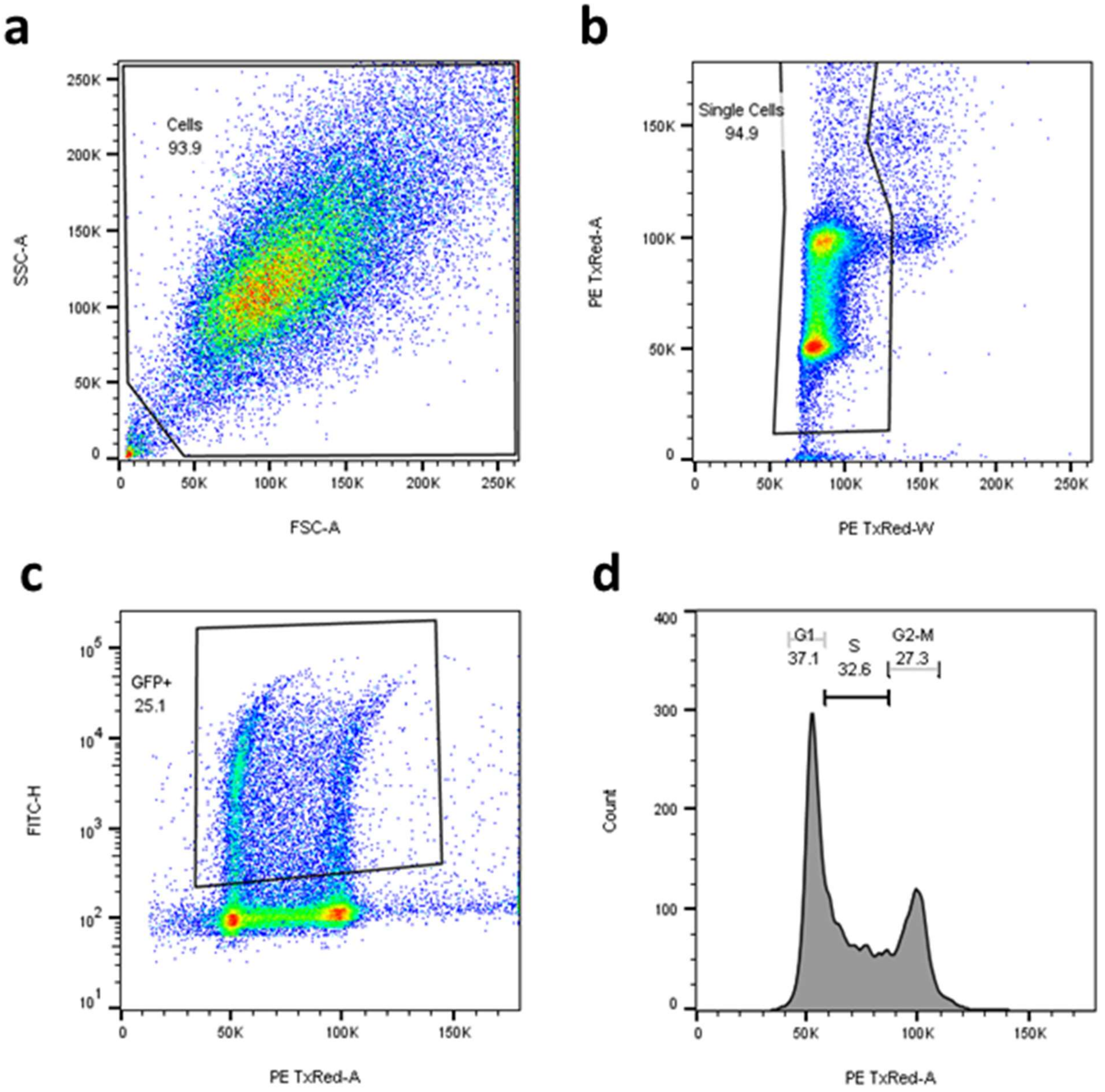

**Supplementary Figure 12**: Flow cytometry gating strategy using sample GFP-BRC8-2 (No IR), as example. **a** First, cells were plotted SSC-A / FSC-A to remove cellular debris. **b** Subsequently, the selected cells were plotted PE TxRED-A / PE TxRED-W (DNA-PI channel) to gate single cells. **c** These single cells were plotted FITC-H (GFP fluorescence channel) / PE TxRED-A (DNA-PI channel) to gate the GFP positive cells which were used to generate the cell cycle profiles similar to the one shown in **d**.
